# Microbial trait multifunctionality drives soil organic matter formation potential

**DOI:** 10.1101/2024.05.24.595733

**Authors:** Emily D. Whalen, A. Stuart Grandy, Kevin M. Geyer, Eric W. Morrison, Serita D. Frey

## Abstract

Soil microbes are a major source of organic residues that accumulate as soil organic matter (SOM), the largest terrestrial reservoir of carbon on Earth. As such, there is growing interest in determining the microbial traits that drive SOM formation and stabilization; however, whether certain microbial traits consistently predict SOM accumulation across different functional pools (e.g., total vs. stable SOM) is unresolved. To address these uncertainties, we incubated individual species of fungi in SOM-free model soils, allowing us to directly relate the physiological, morphological, and biochemical traits of fungi to their SOM formation potentials. We find that the formation of different SOM functional pools is associated with distinct fungal traits, and that ‘multifunctional’ species with intermediate investment across this key grouping of traits (namely, carbon use efficiency, growth rate, turnover rate, and biomass protein and phenol contents) promote SOM formation, functional complexity, and stability. Our results highlight the limitations of categorical trait-based frameworks that describe binary (high/low) trade-offs between microbial traits, instead emphasizing the importance of synergies among microbial traits for the formation of functionally complex SOM.

## Introduction

Microbial metabolism shapes the balance between carbon (C) loss and accrual within the soil organic matter (SOM) pool, the largest actively cycling reservoir of C on Earth^1^. While microbial decomposition and respiration (catabolism) are responsible for a significant proportion of annual carbon dioxide emissions from SOM to the atmosphere, these processes are accompanied by the production of microbial biomass (anabolism), and subsequent deposition of cellular residues and extracellular products (i.e., microbial necromass), thereby providing a pathway for the simultaneous accumulation of SOM^2^. As such, microbes are increasingly recognized for their contributions to SOM alongside direct inputs from plants^3–6^. Contemporary theory recognizes SOM formation and persistence as the balance between microbial catabolic and anabolic processes, further regulated by soil physicochemical properties that restrict decomposer access to substrates (e.g., pore connectivity, aggregation, mineralogy), as well as biotic factors that influence the outcome of microbe-substrate interactions (e.g., microbial traits such as enzymatic repertoire or growth efficiency)^7–10^. Owing to this important role of microbial anabolism in the soil C cycle, a growing number of studies attempt to link microbial traits to SOM formation and stabilization^11–14^; however, whether certain microbial traits consistently predict organic matter accumulation across different SOM pools is unresolved.

Soil organic matter is structurally and molecularly complex, being comprised by multiple functional pools that contribute to the myriad of ecosystem functions provided by soils, including nutrient cycling, C storage, water retention and erosion prevention^15–17^. Such functional pools include particulate organic matter (POM) and mineral-associated organic matter (MAOM). POM is characterized by larger organic matter fragments that are relatively accessible to microbial decomposers, promoting soil C and nutrient cycling, while MAOM is comprised of smaller biopolymers and monomers in close association with silt and clay minerals, rendering this SOM relatively protected from microbial decomposition and promoting C storage in soils^18,19^. POM and MAOM are integrated into the complex three-dimensional structure of soil aggregates, further protecting SOM from microbial decomposition through physical occlusion and modification of micro-scale environmental gradients (e.g., O_2_ availability)^18,20^.

While these SOM pools differ broadly in their relative stabilities, each pool can contain organic matter with a range of turnover times (e.g., “fast cycling” MAOM or occluded POM)^21–23^. Indeed, emerging understanding suggests that mineral sorption should be viewed as a reversible process that temporarily increases C retention in soils^8^ and that greater spatial heterogeneity and molecular diversity of SOM (so-called ‘functional complexity’^17^) promote its retention, while simultaneously maintaining other vital ecosystem functions (e.g., nutrient cycling, water retention)^15–17^. This view underscores the need for a holistic understanding of the role of microbial traits in the formation of SOM functional pools, and in the generation of SOM functional complexity^17^. Because SOM pools are formed via multiple plant and microbial pathways^6,18^, it is likely that their formation is controlled by distinct traits.

Trait-based frameworks adapted from plant ecology^24–26^ are increasingly applied in microbial ecology to distill the compositional and functional complexity of microbial communities, and to link microbial identity to emergent ecosystem functions, including SOM formation^27,28^. Prior frameworks have theorized the role of several microbial trait categories (e.g., physiology, growth morphology and cellular/extracellular biochemistry)^27–30^ in SOM formation and persistence. However, empirical evidence is limited, with a majority of studies measuring a single physiological trait—carbon use efficiency (CUE), the proportion of substrate C that microbes allocate towards growth relative to respiration^31^. These studies have proposed a positive empirical relationship between CUE and soil C^11,14,30,31^, suggesting that microbes that maximize investment in CUE at the expense of other traits (i.e., genetic and/or physiological tradeoffs)^32^ will make the largest contributions to SOM formation^27,28^. However, evidence is mixed^13,14,33,34^, suggesting that current trait-based frameworks emphasizing single traits (e.g., CUE) – or binary tradeoffs between traits – are insufficient to describe the microbial controls on SOM accumulation.

Whereas soil microbial CUE appears to positively correlate with total SOM-C^14,35,36^, its relationship with relatively stable pools of SOM, such as MAOM, may be positive, negative or neutral^13,33,34^, and understanding is limited by the paucity of studies evaluating stable subfractions of MAOM alongside CUE^33^. Emerging evidence suggests that CUE may be decoupled from MAOM accumulation under circumstances where MAOM-C is predominantly plant-rather than microbial-derived (e.g., in temperate forests^34^ or detritusphere soils^37^).

Microbial CUE may also become decoupled from stable SOM formation if other traits strongly influence the incorporation of microbial residues into mineral-associated or otherwise protected pools^12,18^. It was recently hypothesized that microbial physiological traits, such as CUE and growth rate, act as initial filters on the “feedstock”^18^ of microbial residues available for incorporation into SOM functional pools, while additional microbial traits regulate the subset of those residues that become stabilized in soils (i.e., MAOM formation traits)^18^. For example, physiological traits such as CUE may predict total microbial residue inputs to soils (except see^38^), while growth morphology (e.g., fungal hyphal surface area) and cellular and extracellular residue chemistries (e.g., protein contents) are likely to influence the degree and strength of resulting organo-mineral interactions^18,29^. A comprehensive approach that accounts for the groupings of physiological, biochemical and morphological traits of microbes involved in SOM formation would more accurately capture the multidimensional nature of microbial trait expression^39^, allowing for the identification of potential interactions and synergies between microbial traits that may be necessary for the formation of different SOM pools. However, empirical tests of such a framework are currently lacking.

Here, we characterized a suite of microbial physiological, biochemical, and morphological traits hypothesized to be important for the production of microbial feedstock^18^ (i.e., cellular and extracellular residues) and its subsequent incorporation into SOM functional pools, including relatively persistent pools of SOM (i.e., formation traits)^18^. We took a comprehensive approach, accounting for the role of these multidimensional trait profiles in the formation of multiple SOM pools, including total SOM-C, POM, MAOM, water-stable aggregates, and chemically and/or biologically stable fractions of SOM. Our study focused on fungi, which dominate the microbial biomass pool^40,41^, necromass pool^42,43^, and decomposition processes^44,45^ in many ecosystems, and are therefore likely a major source of microbial-derived SOM^29^. We employed a unique study design with individual fungal species incubated axenically in SOM-free model soils, allowing us to directly relate a species’ multidimensional trait profile to its SOM formation potential. Model soils have been successfully employed to monitor microbial formation of SOM from simple C substrates (Pronk et al., 2013, 2017; Del Valle et al., 2022), providing direct evidence that microbes are a source of SOM quantity and chemical complexity (Kallenbach et al., 2016). Importantly, such simplified systems allow direct inferences to be made about relationships between the trait profiles of microbes and the SOM that they generate. We hypothesized that CUE would be an important predictor of total soil C, but that other fungal traits (e.g., variables related to hyphal morphology or biomass chemistry) would emerge as drivers of fungal contributions to relatively stable pools of SOM. Our results partially support our hypothesis, illuminating a key grouping of fungal traits involved in SOM formation, with distinct traits emerging as important predictors of total soil C, MAOM-C and persistent subfractions of the MAOM pool. We propose a holistic framework for understanding the microbial role in SOM formation, identifying synergies among microbial traits that promote SOM quantity, stability and functional complexity.

## Results

### Fungal Traits

We characterized a suite of physiological (CUE, growth rate, turnover rate, extracellular enzyme production), biochemical (biomass chemistry, melanin content), and morphological (hyphal length and surface area per soil volume) traits for eight fungal species spanning three phyla (Basidiomycota, Ascomycota, Mucoromycota [subphylum Mucoromycotina]). To evaluate each species’ performance of multiple traits, we calculated a metric of ‘trait multifunctionality’ by adapting an approach for characterizing ecosystem multifunctionality^46,47^ (details provided in Methods). This metric accounted for both the presence of traits (value > 0) within an isolate’s trait profile and their relative values. While our focus in this experiment was on relationships between fungal trait profiles and the formation of different SOM pools, we present results at the species and phylum levels to aid interpretation and understanding of the range of trait (and SOM pool) values spanned by the fungal isolates included in this study. Relative values are not indicative of innate or immutable characteristics of the fungal isolates, but rather their relative trait or SOM values under the specific experimental conditions of this study. Fungal species exhibited distinct trait profiles (MANOVA: *P* = 0.001; Fig. 1; Fig. S1), corresponding with differences in trait multifunctionality (one-way ANOVA; *P* = 0.003; Fig. 1; Fig. S2). Certain species (e.g., *H. minutispora*, *P. lacerum*) exhibited intermediate values across a wider range of traits (Fig. 1), corresponding with higher trait multifunctionality scores (Fig. 1; Fig. S2), while other species (e.g., *P. stipticus*, *Gymnopus sp.*) exhibited a greater degree of specialization, with high values restricted to a more limited suite of traits (lower trait multifunctionality scores). Correspondingly, these species had relatively low values across the remaining suite of measured traits (Fig. 1; Fig. S2).

**Figure 1.**
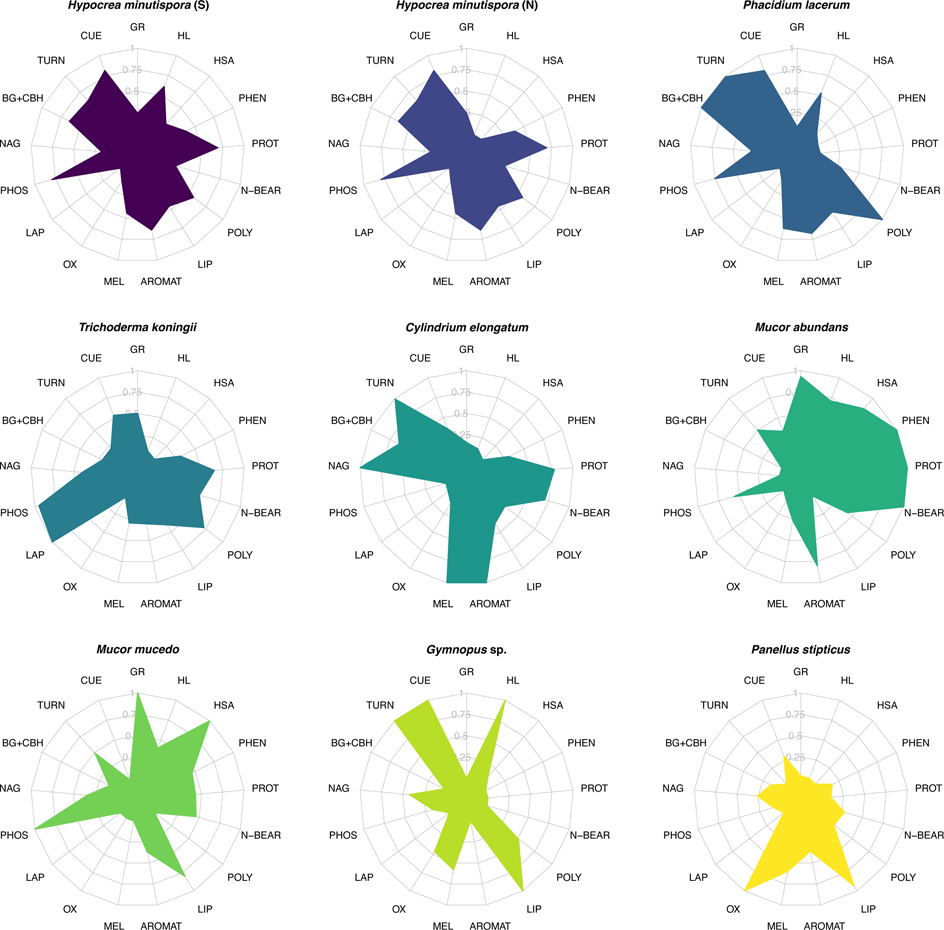
Radar plots illustrating trait profiles of fungal species. Trait values are scaled between 0 to 1 across species for each individual trait measurement, where 0 represents the lowest relative value of trait expression, and 1 represents the highest relative value (n = 4, except *P. stipticus,* n = 3). Species are ordered from high trait multifunctionality scores (top row) to low trait multifunctionality (bottom row), and the color associated with each fungal species is carried throughout subsequent plots. Text abbreviations around the radar plots represent individual traits. Trait abbreviations are as follows: GR = growth rate (average of two metrics); CUE = carbon use efficiency (average of three metrics); MEL = melanin; TURN = biomass turnover (reflected such that a high value corresponds with high turnover rate); BG + CBH = combined activity of beta-glucosidase and cellobiohydrolase; NAG = N-acetyl-glucosaminadase activity; PHOS = acid phosphatase activity; LAP = leucine aminopeptidase activity; OX = combined activity of phenol oxidase (ABTS) and peroxidase (TMB); AROMAT = biomass aromatics; LIP = biomass lipids; POLY = biomass polysaccharides; N-BEAR = biomass N-bearing compounds; PROT = biomass proteins; PHEN = biomass phenols; HSA = hyphal surface area; HL = hyphal length.

Trait profiles exhibited clear groupings at the phylum level (MANOVA: *P* = 0.001; Fig. 2). Higher growth rates were associated with Mucoromycotina and Ascomycota (*P* < 0.001, one-way ANOVA, Fig. S1; post-hoc Tukey HSD results in Table S1), while higher CUE was associated with Basidiomycota and Ascomycota (all *P* < 0.05; Fig. S1; Table S1). Biomass turnover length was generally shorter (i.e., faster turnover rates) among the Ascomycota and one Basidiomycota species (*Gymnopus sp.*; *P* = 0.004; Fig. S1). Biomass protein, phenol, and N-bearing contents were highest among the Mucoromycotina and Ascomycota (all *P* < 0.05; Table S1; Fig. S1). Biomass melanin concentrations were highest among Basidiomycota and Ascomycota species (*P* < 0.001; Table S1), particularly *C. elongatum* and *P. lacerum* (Fig. S1). Mucoromycotina exhibited the highest hyphal surface areas (*P* < 0.001), driven in part by their large relative hyphal diameters (*P* < 0.001; Table S1; Fig. S1). Intermediate to high hyphal lengths were observed for individual species within each phylum (particularly *Gymnopus sp*., *H. minutispora* [sporulating] and *M. abundans*; Fig. S1). Oxidative enzyme (OX) activities were highest among the Basidiomycota (*P* = 0.001), while the hydrolytic enzymes BG and CBH were highest among the Ascomycota (*P* < 0.001; Table S1; Fig. S1). PHOS activity was higher among the Ascomycota and Mucoromycotina isolates than the Basidomycota (*P* = 0.011; Table S1; Fig. S1). The potential activities of LAP and NAG did not differ significantly by phylum (*P* = 0.09 and *P* = 0.29, respectively; Table S1), though there were significant differences at the species-level (*P* < 0.01; Fig. S1).

**Figure 2.**
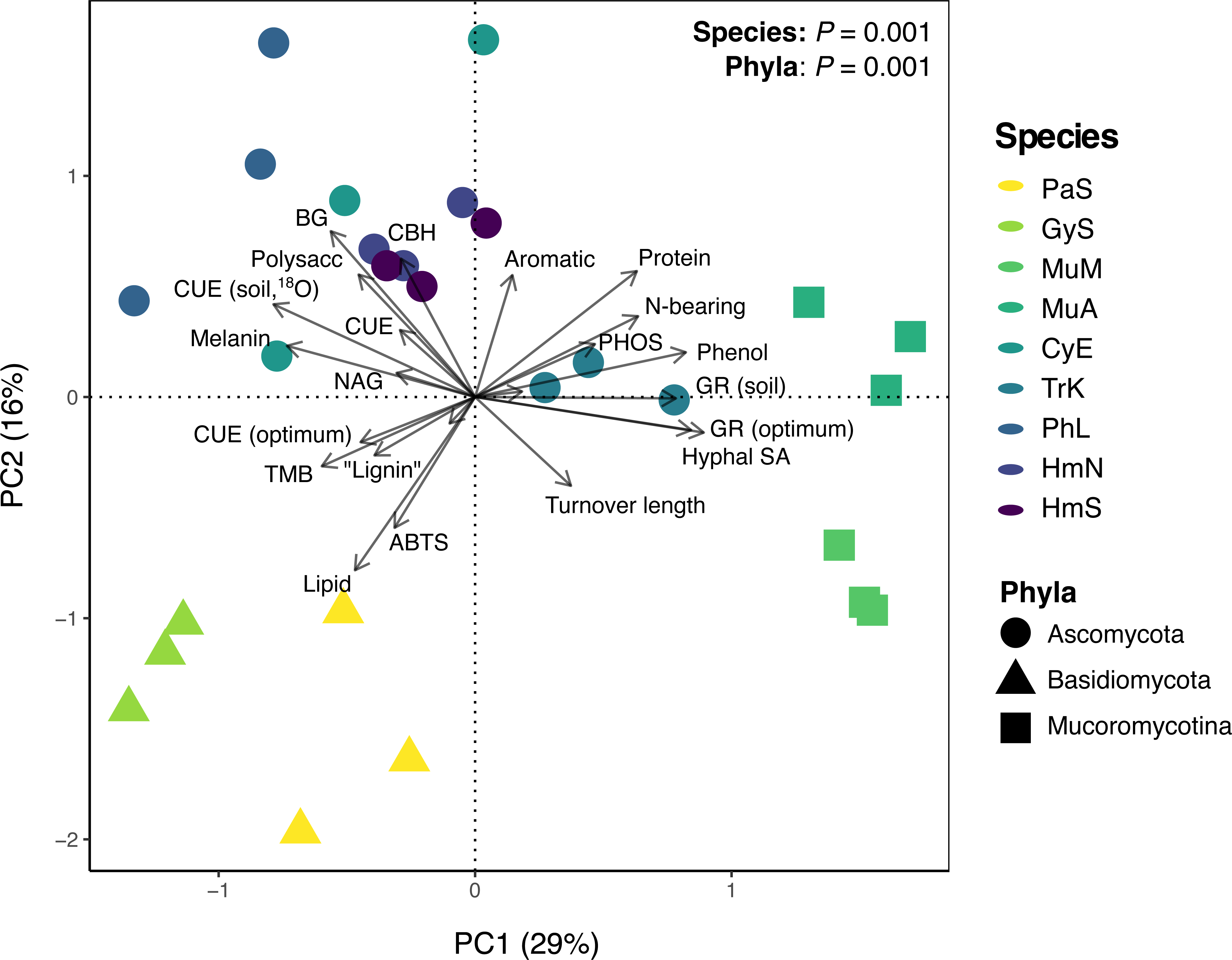
Principal components analysis (PCA) of fungal trait profiles (n=3). The two most explanatory PC axes are presented (PC1, PC2), which collectively explained 45% of variation in fungal trait data. The color of sample points represents fungal species, while point shape represents fungal phyla. Vectors indicate individual trait measurements. The results of separate MANOVA analyses for species and phyla are presented (both *P* = 0.001). Fungal species abbreviations are consistent throughout this manuscript and are as follows: HmS = *H. minutispora* (sporulating); HmN = *H. minutispora* (non-sporulating); PhL = *P. lacerum*; TrK = *T. koningii*; CyE = *C. elongatum*; MuA = *M. abundans*; MuM = *M. mucedo*; GyS = *Gymnopus sp.*; PaS = *P. stipticus*.

### Fungal SOM Formation

Fungi were incubated axenically in initially SOM-free, substrate-amended model soil for between 3-6 months depending on species’ individual growth dynamics. Each species received the same total quantity and rate of substrate-C amendment, while the timing of amendments and harvests were made on a species-specific basis (based on respiration rates, Fig. S3; see Methods for details). Species’ contributions to total soil C, MAOM-C, POM-C, water-stable aggregates, and biologically stable or chemically stable subfractions of SOM were assessed. Fungal contributions to each of the measured functional pools were compared against sterile control soils (model soil mixture + substrate) to account for possible abiotic retention of substrate-C. Fungal species differed greatly in their contributions to different SOM functional pools (Fig. 3; Fig. S4; *P* < 0.05 for all pools). Across all species, between 31-46% of the total substrate-C added over the course of the experiment remained as SOM-C (on average; Fig. S4). The average proportion of total soil C comprised by the MAOM fraction ranged between 63-91% across species, while the average proportion in POM was 9-37% (Fig. S4; all *P* < 0.05). Total soil C varied by species (*P* = 0.007; Fig. 3) and was generally highest among the Ascomycota and Basidiomycota isolates, though these differences were not significant at the phylum-level (*P* = 0.301; Table S2). Contributions to MAOM-C and water-stable aggregates varied by phylum (*P* = 0.045 and *P* < 0.001, respectively; Table S2), with the highest average values observed among Ascomycota and Basidiomycota isolates (post-hoc comparisons: *P* < 0.05; Table S2). Biologically and chemically stable SOM also varied by phylum (both *P* < 0.01; Table S2), with the largest contributions in this case observed among the Ascomycota and Mucoromycotina (post-hoc comparisons: *P* < 0.05; Table S2). We thus sought to understand how fungal contributions to different SOM functional pools were related to their respective trait profiles and functioning.

**Figure 3.**
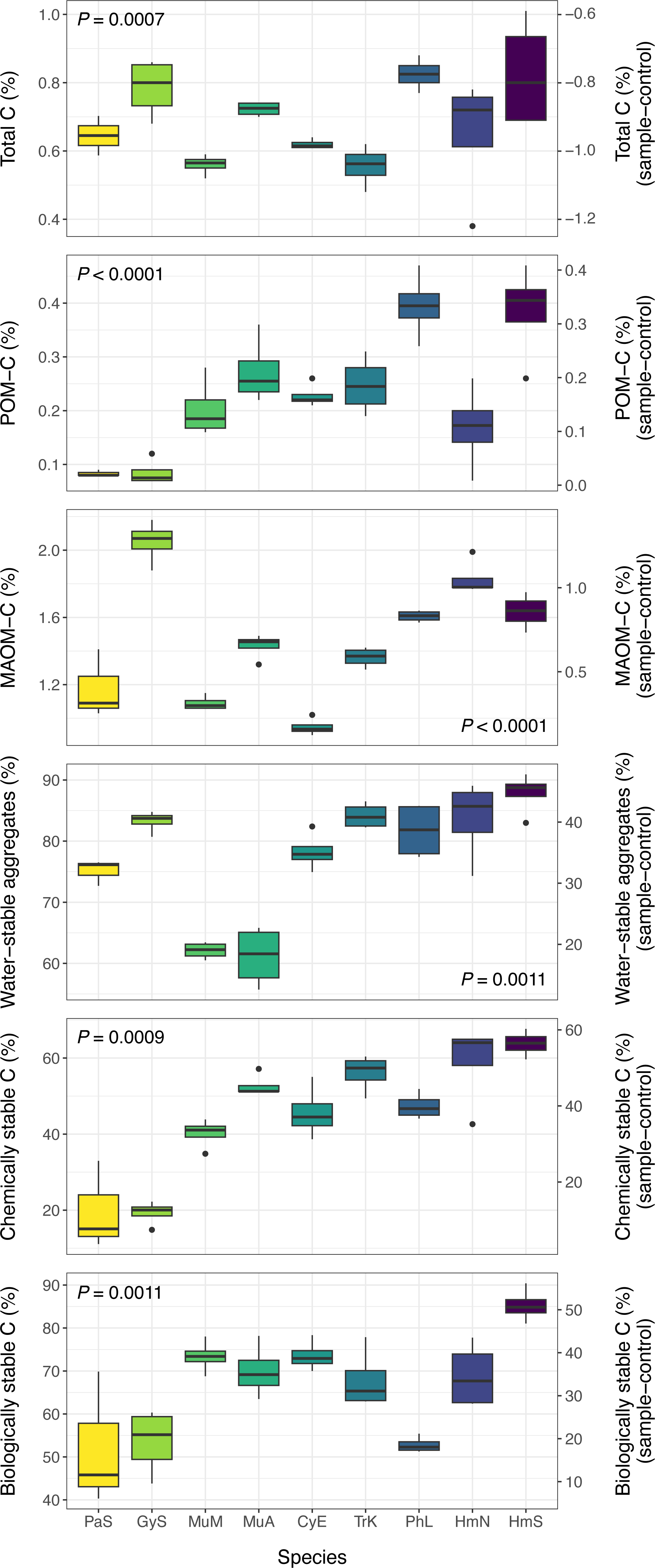
Isolate contributions to SOM functional pools, presented as total uncorrected values (lefthand Y-axis) and corrected values (righthand Y-axis), which were calculated as the difference between inoculated (fungal) and sterile control samples (n = 4, except *P. stipticus*, n = 3). Corrected values for total C are negative, as the sterile control samples contained higher C concentrations than the inoculated samples, due to the absence of respiratory CO_2_ losses. Boxplots represent 25^th^ and 75^th^ percentile, median and outlying points. Results (p-values) of one-way ANOVA or Kruskall-Wallis tests are presented for each SOM functional pool (all *P* < 0.05).

### Traits Predicting SOM Formation

We used partial least squares regression (PLSR) to identify fungal traits associated with the formation of each SOM functional pool. Across our dataset, CUE was one of the most important traits regulating the formation of MAOM and total C (Fig. 4a; strong positive loadings, high VIP scores in PLSR models), whereas biomass protein and phenol contents were the most important traits contributing to the formation of chemically and biologically stable SOM (Fig. 4a). Growth rate was also an influential trait variable in each PLSR model, loading positively on the most explanatory latent factor for biologically stable SOM (and to a lesser extent, chemically stable SOM) versus negatively on the latent factors for total C and MAOM-C. These relationships were reflected in the significant positive correlations between CUE and total C or MAOM-C (Fig. 4b; R^2^ = 0.43-0.67; *P* ≤ 0.0002), and in the positive correlations between biomass protein content and chemically or biologically stable SOM (R^2^ = 0.40; *P* ≤ 0.0004). CUE was also positively correlated with the proportion of water-stable aggregates (R^2^ = 0.65; *P* < 0.0001).

**Figure 4.**
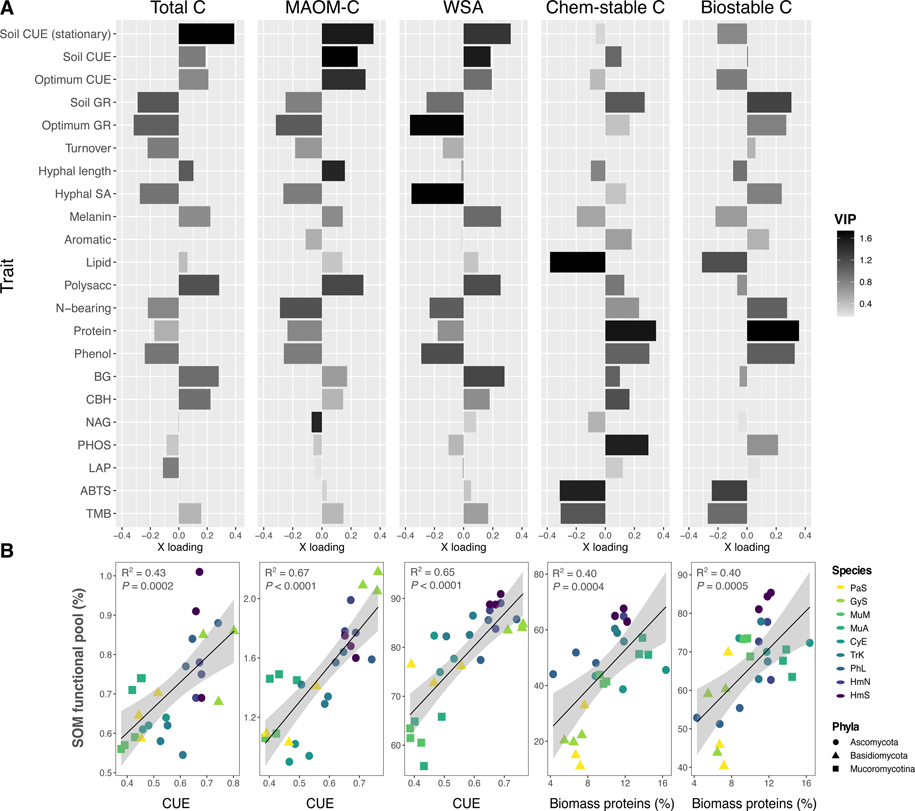
(A) Bar plots of trait loadings on the first (most explanatory) PLSR latent factor for each SOM functional pool, accounting for 50-78% of variation in each pool. From left to right: total C, MAOM-C, water-stable aggregates (WSA), chemically stable C, and biologically stable C. Bar color is shaded to represent variable importance scores (VIP) for each trait variable (n=3), indicating the overall importance of each trait to the PLSR model, integrating latent factor 2, and in some cases, factor 3. Full PLSR model results are reported in Table S3, and a version of this plot with POM included is presented in Fig. S4. Optimum growth rate and CUE were measured in liquid culture. All growth rate and CUE assays were conducted during log-phase, unless specified at stationary growth. (B) Linear regressions (n=27) between the trait variable with the strongest loading on PLSR1 for each SOM functional pool, where regression plots correspond with the functional pool whose PLSR results are presented directly above. For regressions involving CUE, an average CUE value was calculated across the three CUE metrics included in this study (liquid culture log-phase, soil log-phase, soil stationary growth). Biomass proteins represent the relative abundance of proteins in the biomass of fungal isolates grown in liquid culture, as quantified by Py-GC/MS. Point color represents fungal species, as in prior plots, with trait multifunctionality and SOM formation potential generally increasing from light to dark colors (Fig. S2).

Principal components analysis (PCA), used to visualize differences in trait profiles across species and phyla, demonstrated similar patterns of association between fungal traits and SOM pools (Fig. 2). Higher biomass protein, phenol and N-bearing contents, as well as fungal growth rates, loaded positively on PC1 (associated with Mucoromycotina and certain Ascomycota isolates), while CUE loaded negatively on PC1 (associated with the Basidiomycota and remaining Ascomycota isolates). Correspondingly, PC1 was generally associated with greater proportions of chemically and biologically stable C (quadratic polynomial regressions in Fig. S6; R^2^ = 0.32 and 0.28, and both *P* ≤ 0.05), whereas it was negatively correlated with total C and MAOM-C (linear regressions; R^2^ ≈ 0.2 and *P* < 0.05 for both).

The results of these analyses (Fig. 1-4) led us to hypothesize that microbial taxa with intermediate to high performance across this key grouping of traits (namely, CUE, GR and biomass protein and phenol contents) would be most proficient at forming SOM across multiple functional pools. To test this hypothesis, we calculated a metric of ‘SOM formation potential’ to characterize each species’ average contributions to the measured SOM functional pools. The same approach that was used to calculate trait multifunctionality^46,47^ was adapted for this purpose by using SOM pools as input values instead of fungal traits. We found that trait multifunctionality was strongly positively correlated with SOM formation potential (Fig. 5; R^2^ = 0.70; *P* < 0.0001), and this relationship was consistent whether all measured functional pools (Fig. 5) or only the putatively stable pools of SOM were included in the calculation (Fig. S7; R^2^ = 0.65; *P* < 0.0001). To illustrate this point, *Gymnopus sp.* (Basidiomycota) exhibited a high degree of specialization in its trait profile and had a relatively low trait multifunctionality score (0.31; Fig. S2). Correspondingly, *Gymnopus sp.* made large contributions to a small subset of SOM functional pools, namely MAOM-C, but had one of the lowest levels of both chemically and biologically stable C of all the isolates (Fig. 3). In contrast, *H. minutispora* (Ascomycota) exhibited high trait multifunctionality (0.40 for HmS; Fig. S2), including intermediate CUE, growth rate, and biomass protein contents, and formed intermediate to high levels of SOM across all measured functional pools. Overall, the five fungal species with the highest SOM formation potentials (Fig. 5; Fig. S2) were associated with intermediate to high trait values (>0.5 when species’ trait values were scaled 0 to 1) for CUE, turnover rates, and PHOS activity (Fig. 1). Four out of the five taxa were associated with intermediate to high values for biomass proteins, phenols, polysaccharides and aromatics.

**Figure 5.**
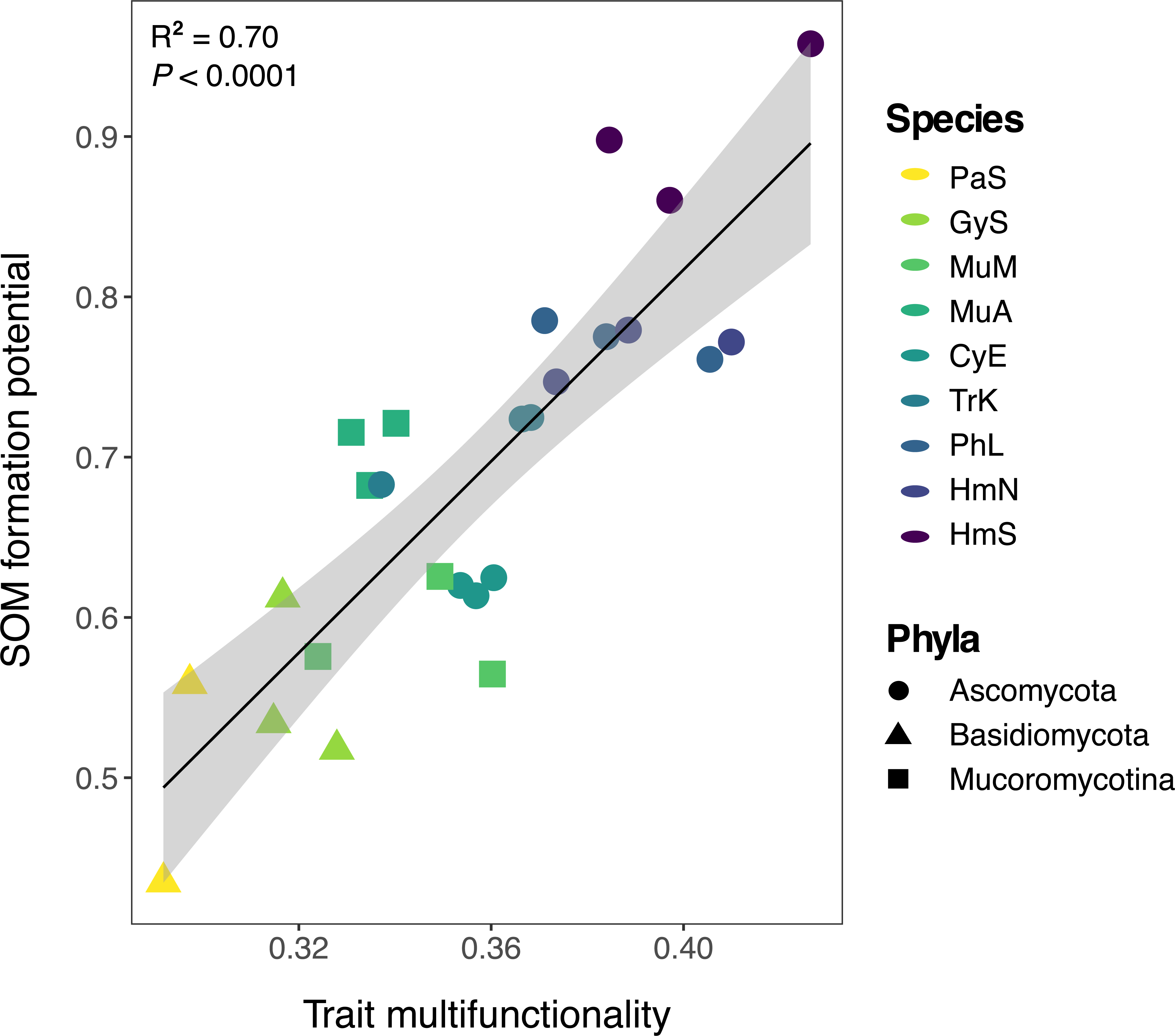
Linear regression between trait multifunctionality and SOM formation potential (n=27). Point color represents fungal species, while point shape represents fungal phyla. The SOM pools included in the calculation of average SOM formation potential included: total C, POM, MAOM, water-stable aggregates, chemically stable C and biologically stable C. One PaS replicate with significantly lower trait multifunctionality (∼0.2) is removed from the plot for ease of visualizing differences across other samples, but does not change the interpretation of the results (both *P* < 0.0001). ANOVA results are presented for the model that included all replicates (R^2^ = 0.70; *P* < 0.0001).

### Chemistry of Fungal-Derived SOM

In addition to evaluating fungal contributions to SOM quantity, we sought to understand whether species identity influenced SOM chemistry. We had two main objectives: (1) evaluate whether fungal production of specific compounds is related to the stability of SOM produced; and (2) characterize the molecular diversity of SOM generated by each fungal species^17^. Using a ramped pyrolysis gas chromatography/mass spectrometry (Py-GC/MS) approach, we found that fungal species generated SOM (Fig. 6a) and MAOM (Fig. S8) with distinct chemistries (both *P* ≤ 0.001), and this species-level variation was apparent within each pyrolysis thermal fraction (Fig. S9; PERMANOVA, all *P* ≤ 0.001). In the ramped pyrolysis approach, organic compounds that resist pyrolysis at lower temperatures, but combust at higher temperatures, are assumed to have higher thermal stabilities, and thus to comprise relatively persistent pools of SOM^48^. In general, the relative abundances of proteins, N-bearing compounds and phenols increased as pyrolysis temperature increased (abundances peaked between 444-600°C), suggesting that these fungal-derived compounds were relatively stable in soils (Fig. S10). In contrast, the relative abundances of polysaccharides and lipids peaked between 330-396°C and generally declined at higher pyrolysis temperatures, suggesting that these compounds were less thermally stable. Consistent with these results, we found that the average relative abundances of phenols and proteins in soils at the end of the long-term incubation were positively correlated with the proportion of chemically stable SOM produced by each species (Fig. 6b-c; R^2^ = 0.57 and 0.33, respectively; both *P* ≤ 0.0003). Additionally, fungi with relatively high SOM formation potentials tended to produce thermally stable SOM characterized by a higher relative abundance of proteins and phenols (HmS, HmN, PhL, TrK, MuA in Fig. S10). These fungi also generally produced SOM that was more chemically diverse, and we observed a positive correlation between the chemical diversity (Shannon index) of thermally stable SOM (735°C thermal fraction) and the proportion of chemically stable C formed by each species (Fig. S11; R^2^=0.55; *P* < 0.0001).

**Figure 6.**
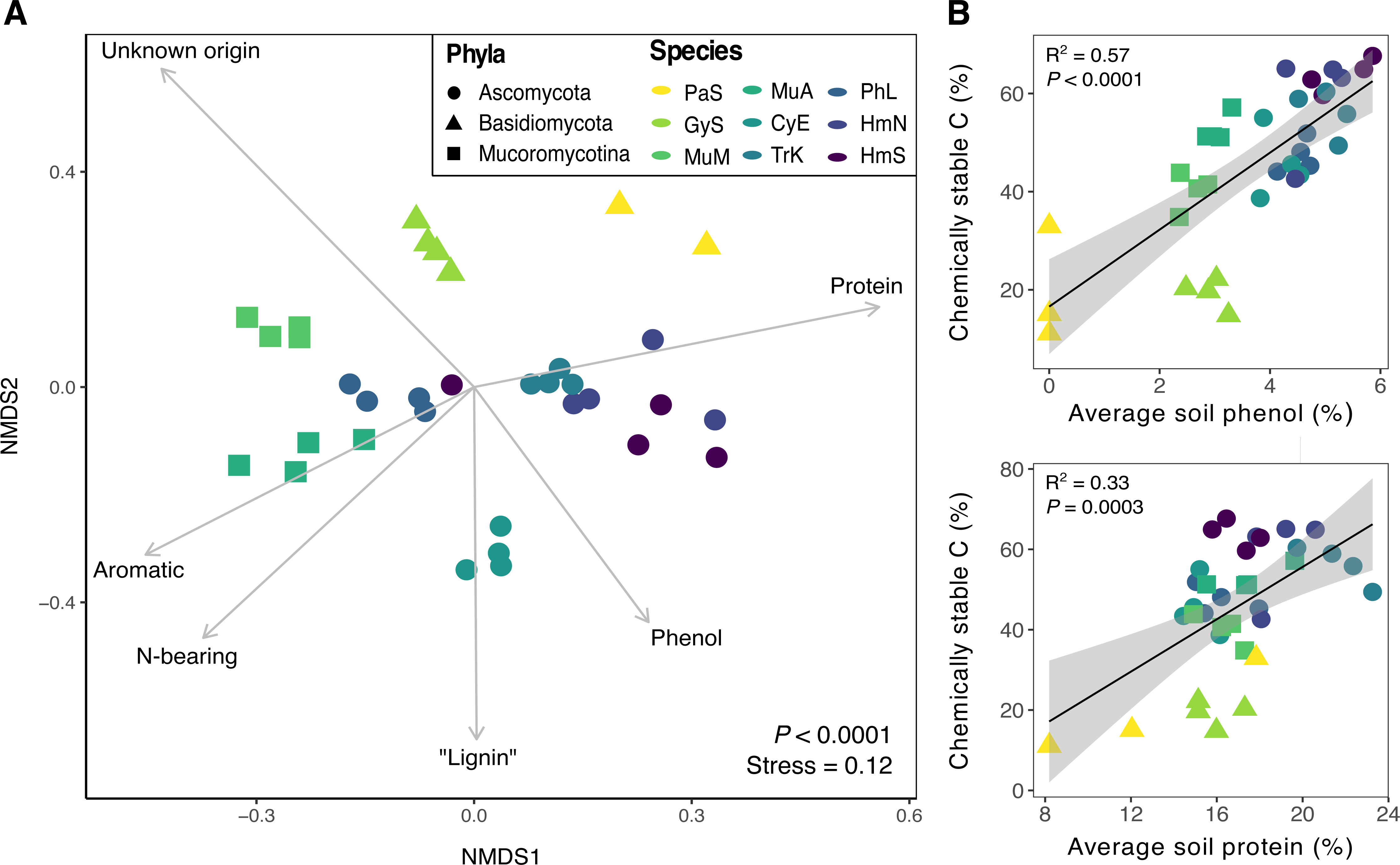
(A) NMDS ordination of SOM chemical composition (individual compound-level data) after long-term incubation with fungal isolates (n = 4, except *P. stipticus,* n = 3). Results for a single temperature fraction (503°C) from the ramped pyrolysis GC/MS analysis are presented here for simplicity. Significant variation in species’ SOM chemistries were observed for each thermal fraction (all *P* < 0.001; PERMANOVA), and full results are shown in Fig. S8. Point color represents fungal species, while point shape represents fungal phyla. Environmental vectors representing broad chemical compound classes were included in the plot if significant (*P* < 0.05). (B) Correlations between the relative abundances of phenols (top) or proteins (bottom) in soils at the end of the long-term incubations and the proportion of chemically stable C produced by each fungal species (n=27).

## Discussion

Fungi differed considerably in their formation and stabilization of SOM based on variation in their trait profiles, with distinct fungal traits emerging as important controls on the formation of different SOM functional pools. Total SOM and MAOM formation were primarily associated with higher CUE, whereas the formation of chemically and biologically stable SOM was associated with higher fungal growth rates and biomass protein and phenol contents. Species with higher trait multifunctionality, especially those with intermediate to high performance across this key grouping of traits, exhibited the greatest capacities for SOM formation across functional pools. We present a holistic framework for understanding the role of fungal traits in SOM formation, proposing that synergies among these traits promote SOM quantity, stability and functional complexity^17^.

Similar to previous experiments employing model soils to investigate SOM formation, we demonstrate that microbes can form SOM that is chemically and functionally diverse, absent complex plant inputs^14,33^. Additionally, we provide direct evidence that fungi contribute to SOM formation and stabilization via multiple pathways within soils, from aggregate formation and particulate hyphal residue inputs within the POM (>53 μm) fraction, to the formation of relatively stable fractions of mineral-associated SOM. Prior frameworks have emphasized the role of fungi in soil particle arrangement and enmeshment, suggesting that fungi are particularly central to the formation of soil macroaggregates^49–51^. However, these frameworks have been less clear on the potential role of fungi as agents of microaggregate and stable MAOM formation, often suggesting that bacteria are primarily responsible for the decomposition of hyphal residues and simultaneous production of biofilms (including EPS and other extracellular compounds) that become major inputs to MAOM^29,52–54^. Here, we demonstrate that in a bacteria-free environment, fungi generated chemically and functionally diverse SOM and MAOM, which included fractions that resisted chemical and biological destabilization. While hyphal decomposition and EPS production by bacteria surely play important roles in MAOM formation in natural soils, our results emphasize that the direct contributions of fungi should not be overlooked. Fungal inputs may include cell wall fragments, cytosolic components from lysed cells, EPS originating from hyphal coatings, as well as other extracellular metabolites (e.g., organic acids, sugars)^29,55–57^. We found that fungal species differed in their contributions to SOM functional pools – likely because of variation in these different pathways of fungal-derived SOM formation – and differences in SOM formation potential were linked to variation in fungal traits.

We observed a pattern among our study species consistent with the theory of ecological trade-offs among microbial generalists versus specialists^58,59^; here, we adapt this theory in the context of microbial traits. Certain species exhibited high trait values within a limited suite of traits (“functional specialists”) whereas other species exhibited intermediate values across a wide range of traits (“functional generalists”), which corresponded with higher trait multifunctionality scores. We found that a species’ trait multifunctionality was an important predictor of its SOM formation potential, suggesting that generalists who invest at intermediate levels across a wider range of trait categories (i.e., “jack of all trades”) are more likely to have the genetic and/or physiological capacity for traits important to SOM formation, even if they do not exhibit maximal performance of any given trait. This is likely because different fungal traits were associated with the formation of each SOM functional pool, and certain fungal traits may be positively correlated with one functional pool but neutrally or negatively associated with another. Thus, species with “bundles of functions with synergies”^60^ may be most likely to possess the capacity for SOM formation across multiple functional pools. While microbes exhibit fundamental tradeoffs in genetic and/or physiological investment in different traits^32^, prior frameworks have presented a two-dimensional or binary view of “high” versus “low” investment^27,28^, proposing community-level syndromes associated with soil C cycling (e.g., ‘high yield’ vs. ‘resource acquisition’ vs. ‘stress tolerance’)^28^. Our results suggest that this binary view overemphasizes microbes performing at extreme ends of the trait value continuum^39^, obscuring the importance of microbes with intermediate investment across a wider range of trait categories involved in SOM formation.

Microbial community CUE has been the focus of most empirical studies evaluating the relationships between microbial traits and SOM formation, with evidence generally supporting a positive correlation between CUE and total soil C storage^11,13,14,35^. However, the relationship between CUE and relatively persistent SOM is less clear, with some lab-based studies showing a positive empirical relationship between these two variables^33^ and emerging field-based evidence demonstrating that microbial CUE can become decoupled from stable SOM formation (e.g., in temperate forests)^34^. Our results suggest that intermediate to high growth rates and biomass protein, phenol and N-bearing contents are stronger predictors of fungal contributions to persistent SOM than CUE, but that when fungi possess intermediate CUE alongside these other traits, SOM formation potential is amplified. Multiple species with high SOM formation potential also exhibited intermediate to high biomass turnover rates. Microbial CUE, growth and turnover rates may act as initial filters on the “feedstock” of microbial residues available for incorporation into SOM functional pools, while additional microbial traits (“formation traits,” e.g., cellular and extracellular residue chemistries) regulate the subset of those residues that becomes stabilized in soils^18^. Our results suggest that this specific combination of traits is synergistic for the formation of SOM. Thus, individual species with the capacity to perform these traits (high trait multifunctionality), or communities of microorganisms where these key traits are well-represented, may possess the greatest SOM formation potentials.

An understanding of microbial growth and turnover rates alongside CUE provides a more complete picture of microbial physiology, giving insight into the amount of biomass and cellular necromass inputs to soils per unit time^18,61^. While high CUE has been the focus of discussions around a ‘high yield’ strategy among microbes^28^, CUE is typically defined per unit substrate^62^ and does not guarantee high biomass and necromass yield per unit time. In ecosystems or spatial microsites in soils where C inputs are relatively continuous, it is likely that the integrative effects of microbial growth rates, turnover rates and CUE regulate SOM-C accrual, especially microbial-derived SOM^11,13,61^. Higher growth rates and activity levels among microbes may be associated with faster biomass turnover and more extensive processing of necromass into lower molecular weight compounds, producing “feedstock” for stable MAOM formation^18,63^. Indeed, we observed greater SOM formation potential among species with a higher capacity for the production of hydrolytic enzymes involved in necromass decomposition (e.g., PHOS, BG, CBH), consistent with the idea that microbial processing of particulate OM can increase the reactivity and diffusion of these fragmented residues to soil minerals^8^.

In contrast, if fungi are efficient (i.e., high CUE) but their growth rate is slow, they may promote C retention in soils by limiting the amount of soil C lost via respiration, but this C will not necessarily be converted into stable forms. In field soils, microbial communities characterized by slower growing species may be associated with more direct stabilization of plant-derived, as opposed to microbial-derived SOM^34^. In our study, a Basidiomycota species (*Gymnopus sp.*) had the highest MAOM-C concentrations and one of the highest total soil C concentrations; however, only a small fraction of this C was biologically or chemically stable. Although incubation length was longest for this species (6 mo. compared with 3 mo. for the fastest growing taxon), this slow-growing fungus may have had sufficient access to substrate C throughout the course of the incubation, reducing its need to mine C and nutrients from existing necromass. Overall, our results suggest that more active fungal species with intermediate to high growth rates, CUE, turnover rates, and enzyme activities produce more residues with the potential for incorporation into stable SOM pools compared to taxa characterized exclusively by high CUE.

While physiological traits largely control the pool of available microbial residues, additional microbial traits influence the likelihood that these residues are incorporated into stable SOM pools (“formation traits”)^18^ alongside characteristics of the soil matrix^64,65^. Our results suggest that biomass chemistry, namely protein and phenol contents, may be one such trait influencing the formation of persistent SOM. While the chemical complexity (‘recalcitrance’) of plant and microbial residues is no longer thought to be a dominant control on their long-term retention in soils^9,10^, the sorptive affinity of individual compounds is an important factor influencing organo-mineral associations^8,66^. Proteins comprise an important fraction of MAOM because of their unique surface reactivity (e.g., abundance of polar and charged functional groups)^67,68^ and conformational changes in structure (“unfolding”) upon association with certain minerals^66^. Similarly, N-rich microbial metabolites and amino acids have been shown to preferentially accumulate in MAOM fractions^53,69^ and amino acids are strongly sorbed to Fe and Al (hydr)oxides^70^. Phenols also exhibit strong sorptive affinity for Fe and Al (hydr)oxides and short-range order minerals^71^ and while their presence in field soils is commonly attributed to plant origins (e.g., lignin)^52,72^, the source of many soil phenols is unknown, and a larger fraction may be microbial-derived than is currently recognized, as proposed by Whalen et al.^6^ (e.g., melanin-derived phenols, phenolic microbial metabolites)^14,57^.

We observed a positive correlation between the relative abundances of fungal-derived soil proteins and phenols and the amount of chemically stable SOM produced by fungi. Consistent with a recent field study^48^, these compounds also comprised an important fraction of thermally stable fungal SOM. Since fungal species generated SOM with distinct chemical fingerprints, these results suggest that microbes who produce and release higher quantities of proteins and phenols have a greater potential for the formation of persistent SOM. This finding builds on emerging evidence that microbial community composition influences the chemical composition and stability of SOM^14,33^. Taxa with the highest SOM formation potentials also generated SOM that was more chemically diverse, and the chemical diversity of SOM was positively correlated with its stability. While our data are currently limited to eight fungal species, these results provide preliminary support for the functional complexity model proposed by Lehmann et al.^17^, wherein SOM persistence is a function of its molecular diversity and spatial heterogeneity, which likely increases with the formation of multiple SOM functional pools. Thus, microbial communities and their associated traits may play a role in the emergence of SOM functional complexity in soils, as is recognized for other characteristics of the soil habitat^73^.

Together, our results build on emerging evidence^34,37^ that the formation of different SOM functional pools is associated with distinct microbial traits. We demonstrate that SOM functional complexity and stability is promoted by more ‘multifunctional’ microbes with intermediate to high performance across a key grouping of traits (namely, CUE, growth rate, turnover rate, and biomass protein and phenol contents). Greater structural and molecular complexity of SOM formed by such taxa may promote soil C storage, while also maintaining the vital ecosystem services provided by soil aggregates and actively cycling SOM pools^15–17^. While past studies have focused on individual microbial traits (e.g., CUE), suggesting that microbes that maximize their CUE at the expense of other traits (i.e., ‘high yield’ strategy)^28^ will exhibit the greatest potential for SOM formation, our results emphasize the importance of synergies among microbial traits that promote SOM formation and stabilization across multiple functional pools. Specifically, we demonstrate that CUE alone is inadequate to predict the formation of SOM across all functional pools and may be decoupled from SOM stabilization when formation traits^18^ (e.g., biomass protein and phenol contents) influence the subset of microbial residues that are incorporated into stable SOM pools. In field soils, microbial traits (including both “feedstock” and “formation” traits) may become decoupled from stable SOM formation when the direct sorption of plant compounds represents an important pathway of MAOM accumulation^34^ or when properties of the soil mineral matrix select for the retention of particular compounds^64,65^. Our results demonstrate the importance of accounting for the formation of multiple SOM pools and the synergies among microbial traits that promote their formation. Future research would benefit from a more comprehensive approach that can account for these complexities, given divergent relationships between specific microbial traits and SOM pools.

## Methods

### Selection and culturing of fungal isolates

Eight saprotrophic fungal isolates spanning three phyla (Ascomycota, Basidiomycota, Mucoromycota) were selected for inclusion in our study. The Ascomycota included *Phacidium lacerum* (shortened in figure labels to ‘PhL’), *Cylindrium elongatum* (CyE), *Trichoderma koningii* (TrK), and *Hypocrea minutispora* (Hm). The Basidiomycota included *Panellus stipticus* (PaS) and a *Gymnopus sp.* (GyS), and the two Mucoromycota (subphylum Mucoromycotina) isolates were *Mucor mucedo* (MuM) and *Mucor abundans* (MuA). Fungal cultures were isolated from decomposing leaf litter, soil or sporocarps in control plots at the Harvard Forest Long-term Ecological Research (LTER) site (Petersham, MA, USA), as described by van Diepen et al.^74^. Species identification was confirmed using the Basic Local Alignment Search Tool (BLAST)^75^ and comparison to respective ITS sequences in the NCBI nucleotide database. Taxa were assigned provisional species names as in Morrison et al.^32^. The eight isolates were selected based on three central criteria: (1) they represent taxa that are highly abundant soil and litter saprotrophs in a model northern hardwood forest, the Harvard Forest (based on past meta-barcoding studies)^32^, (2) they represent a broad phylogenetic range and were confirmed in a previous liquid culture study to show substantial variation in key physiological traits (e.g., CUE, growth rate)^32^, and (3) their genomes have been sequenced^32^. The selected isolates were plated from long-term storage slants onto potato dextrose agar (PDA) plates and maintained at 4°C for use in subsequent incubations. Prior to each incubation experiment, a fungal plug was transferred to PDA working plates and grown at 25°C to acclimate to incubation conditions.

### Model soil preparation

Model soil was created from a mixture of organic matter-free sand (Quikrete Premium Play Sand; sieved < 2 mm), silt (Lab-Aids Pure Silt Soil, 500cc) and montmorillonite clay (Sigma-Aldrich K10 Montmorillonite Powder). To remove native organic matter and adjust soil pH, the sand fraction was acid washed in 1% HCl, and the silt fraction was muffled at 550°C for three hours and subsequently treated four times with concentrated 6M HCl. The clay fraction was soaked in a bath of 0.1M CaCl_2_ to make it homoionic. Soil size separates were combined 70:20:10 (sand:silt:clay) by weight to create a sandy loam texture approximating the soil texture at the Harvard Forest LTER, where fungi were isolated. The final pH of the soil mixture was confirmed with a pH probe (pH = 4.8 ± 0.01), approaching the pH of A horizon soils at Harvard Forest (∼4.6 in control plots)^76^. Achieving a lower pH was challenging due to the high buffering capacity of the silt fraction. Once homogenized, 50 g soil was weighed into specimen cups and sterilized by autoclaving three times at 120°C with 24 h rest periods in between each cycle.

### Long-term incubation experiment

To quantify each isolate’s potential to form SOM across different functional pools, each of the eight isolates was incubated axenically in model soils. The moisture content of sterilized soils (50 g) was adjusted to 50% water holding capacity with a sterile liquid media solution (pH = 4.5). Substrate media contained 12.6 g L^−1^ D-glucose (0.42 M C), 1.4 g L^−1^ potato infusion extract (46.6 mM C, 3.8 mM N), 8.53 g L^−1^ MES monohydrate buffer (40 mM), and 0.78 g L^−1^ ammonium nitrate (19.5 mM N). Glucose and potato infusion extract supplied 90% and 10% of substrate C, respectively, and substrate additions were made at a rate of 5.6 mg C g^−1^ soil (soil C:N = 20). Potato infusion extract was used to provide essential micronutrients and cofactors required for fungal growth. While other C-supplying substrates were considered, glucose was selected due to its rapid assimilation and low level of discrimination among microbial taxa, including saprotrophic fungi^77–79^. A detailed rationale for this decision is provided in the supplement (see ‘Key experimental design decisions & rationale’). Soils were inoculated under sterile conditions with a 5 mm diameter agar plug (50 ml PDA/plate) of the respective fungal culture and sealed in autoclaved quart-size (∼946 ml) mason jars with lids containing rubber septa. Jar flushing (O_2_ replenishment) and gas sample collection (respiration measurements) were conducted using sterile needles and syringes fitted with 0.2 μm filters (PES membrane; Corning).

Soils were incubated at 25°C for between 3-6 mo., depending on the growth dynamics of each fungal species. To provide C and nutrients to sustain fungal growth, two additional substrate amendments (1 ml; 5.6 mg C g^−1^ soil) were applied over the course of the incubation using a sterilized glass syringe and 8-inch stainless steel needle. While more frequent, less concentrated C amendments would have more realistically simulated root exudation rates into soils^14^, a central constraint of our study was the need to maintain axenic conditions of fungal cultures; therefore, we minimized the number of substrate additions to limit the risk of contamination. A total of 840 mg C was added to each 50 g soil unit over the course of the incubation experiment (16.8 mg C g^−1^ soil, prior to respiratory losses). Respiration was monitored every 12-96 hours (depending on isolate growth stage) using an infrared gas analyzer (LI-COR 6252 IRGA), and the timing of substrate amendments was determined on an isolate-specific basis after each isolate had undergone an exponential growth phase (after prior substrate amendment) followed by a period of stationary growth (Fig. S3). The second substrate addition was made once isolate respiration rate had reached ≤ 1 μg CO_2_-C g^−1^ soil h^−1^. Before the third substrate addition, we intentionally induced a period of C and nutrient limitation by maintaining isolates at stationary growth for ∼20 days to facilitate biomass and necromass recycling. Soils were harvested once isolate respiration rates returned to < 1 μg CO_2_-C g^−1^ soil h^−1^ following the third substrate addition. While alternative approaches to standardization were considered (e.g., standardized by time), it was ultimately determined that the selected approach would allow for the most accurate assessment of the relationships between isolate trait profiles and SOM formation (detailed rationale provided in supplementary methods).

Experimental units were monitored regularly for contamination (visually and via respiration measurements), and contaminated samples were promptly removed from the experiment. For all but one taxon (*Panellus stipticus*; Basidiomycota), we achieved at least four replicates that met the criteria for inclusion in subsequent analyses; for *P. stipticus*, we were only able to include three replicates in subsequent analyses due to contamination challenges with this slow-growing taxa (additional details provided in supplementary methods). During incubation, approximately half of the *Hypocrea minutispora* replicates began producing ascocarps and sporulating. We expected that this could influence SOM formation dynamics and alter the chemistry of SOM produced, and thus, samples for this taxon were separately labeled as “sporulating” (HmS) or “non-sporulating” (HmN) (four replicates each). Four sterile control samples (50 g) were established with the equivalent amount of substrate added to each sample containing fungal cultures (840 mg C total) and were harvested 24 hours later to preclude the possibility of contamination. After harvest, soils were gently homogenized by passing them through a 4 mm sieve and subsequently air-dried for 72 hours. Separate subsamples were used for characterization of SOM functional pools, SOM chemistry and a subset of the fungal trait measurements (hyphal lengths, hyphal surface area).

### Trait measurements

A suite of trait measurements was conducted to characterize physiological, morphological and biochemical traits of the fungal isolates. At the end of the long-term incubations, soil was subsampled (3 g) to directly characterize the length and diameter of hyphae produced by each isolate in the incubated soils. Fungal hyphae were extracted from soils and stained following standard methods outlined in Brundrett et al.^80^. Stained hyphae were evaluated at 200x magnification, and the line intersect method^81^ was used to estimate hyphal lengths. Hyphal diameters were assessed at 400x magnification and quantified using AmScope software (AmScope, Irvine, CA). Ten measurements of hyphal diameter were taken and then averaged for each replicate (four replicates per isolate). Hyphal surface area was calculated as follows: HSA = (2 π r h) + (2 π r), where r represents the hyphal radius (1/2 hyphal diameter), and h represents hyphal length. It should be noted that the hyphal length and HSA measurements represent the low end of possible values for these variables, as some of the hyphae produced earlier in the experiment may have been recycled prior to incubation harvest.

For all remaining trait measurements, separate short-term incubations were conducted for each isolate in model soils, with a small subset of measurements conducted in liquid culture as required (3-4 replicates per isolate per trait measurement). All samples were analyzed immediately following incubation harvest. Carbon use efficiency (CUE) and mass-specific growth rates were assessed in both liquid culture and in model soil. We include these multiple metrics to better represent the long-term growth dynamics of fungal taxa during incubation. The liquid culture measurements were conducted in a previous study^32^ and data are included here as a metric of species’ optimum CUE and growth rate under experimental conditions (25°C, C:N = 20; same media solution). Fungal biomass was collected through vacuum filtration and weighed to assess growth, while respiration was monitored using the LI-COR infrared gas analyzer. Mass-specific growth rate was calculated as biomass-C produced biomass-C^−1^ hr^−1^. In a small lab incubation mirroring this liquid culture approach^32^, soil CUE and growth rate were assessed during log-phase growth. Twenty-one experimental units (10 g soil) per isolate were incubated at 25°C for ∼1 week to 1 month (depending on isolate growth rate). Experimental units were destructively harvested at seven time points across each isolate’s respective log-phase growth curve (3 replicates x 7 sampling times), and a ∼0.3 g soil sample was flash frozen and stored at −80°C for downstream DNA extraction. Respiration was measured 12 hours before, and then again immediately before each harvest to assess respiratory losses of CO_2_ (LI-COR analysis, described above). Soil DNA was extracted as a metric of fungal biomass growth using a modified phenol-chloroform extraction procedure designed for clay- and iron oxide-rich soils (*sensu* Whitman et al.^82^; see supplementary methods for full details). DNA concentrations were quantified with a Qubit 3.0 Fluorometer (Life Technologies, Grand Island, NY). Microbial biomass C (MBC) was estimated from DNA concentrations using a published conversion factor of 5.0 to convert μg DNA g^−1^ soil to μg MBC g^−1^ soil^83^. To assess the validity of this approach for our samples, we generated a species-specific conversion factor for each isolate by extracting DNA from a known quantity (1-2 g) of fungal biomass^32^. Because of low replication, we calculated an average conversion factor for all eight isolates (∼4.5), which was comparable to the published conversion factor^83^. After conversion from DNA to MBC, we regressed MBC over time (separately for each isolate) and took the slope of the linear portion of each growth curve to estimate isolate growth rates. Carbon use efficiency was subsequently calculated as (growth/growth + respiration), where respiration rates were acquired by taking the slope of the linear portion of the curve for respiration (μg CO_2_-C produced g^−1^ soil h^−1^) regressed over time, as in Morrison et al.^32^. Standard errors of CUE were calculated by pooling the standard errors of the slopes for growth and respiration^32^.

Soil CUE was also assessed during stationary phase growth using the ^18^O-water tracing method, which quantifies the incorporation of ^18^O-labeled water into DNA to measure gross growth^62,84^. Four additional replicates per isolate (10 g soil) were incubated at 25°C and harvested when each isolate was determined to have reached stationary growth (steady state; based on respiration measurements). Steady state was selected as the timepoint for harvest and for initiation of the ^18^O-water tracing assay to meet the assumptions for estimating biomass turnover. The ^18^O-water tracing assay was conducted following methods recommended in Geyer et al.^62^. Briefly, enriched (∼97 at%) ^18^O-water was diluted with unlabeled filtered deionized water (FDIW) to achieve a target soil moisture enrichment of 20 atom%. Approximately 0.3 g soil was weighed into microcentrifuge tubes, receiving either a labeled (^18^O) or unlabeled (^16^O; control) FDIW solution. Soils were sealed in 50 ml amber vials fitted with rubber septa and flushed with CO_2_-free air. Soils were incubated for 48 hours, whereafter respiration measurements were taken and soils were flash frozen and stored at −80°C until DNA extraction. DNA was extracted following the modified phenol-chloroform protocol^82^ (SI methods). DNA extracts were mixed with a diluted salmon DNA solution (20 μg ul^−1^) to bring total oxygen mass within detection limits and dried overnight in silver encapsulation tins at 60°C. Tins containing dried DNA extracts were sent to the Cornell Stable Isotope Laboratory for 8^18^O quantification.

Total microbial growth (production of new ^18^O-labeled DNA) was estimated using a two-pool mixing model^62,85^. Typically, microbial growth values are scaled to MBC using a ratio of MBC:DNA generated through chloroform fumigation extraction (CFE). Our attempts to extract and quantify biomass from model soils using the CFE approach were unsuccessful, potentially due to the high potential for cell lysates to re-sorb to clean mineral surfaces in these constructed soils. Therefore, we quantified DNA concentrations by Qubit and used the published conversion factor for DNA to MBC as described above^83^. Biomass turnover length and CUE were calculated as in Geyer et al.^62^ For one Basidiomycota species (*Gymnopus sp.*), turnover rate may be artificially inflated relative to other taxa due to especially slow linear-phase growth which may have violated assumptions of the ^18^O method; however, we assume that samples for most species generally met the steady state requirement.

Soils were assayed for potential activities of hydrolytic and oxidative enzymes following standard methods^86,87^. Four replicates (10 g soil) were established for each isolate alongside the experimental units for measuring soil CUE, and were destructively harvested and homogenized before subsampling for enzyme assays. The hydrolytic enzymes cellobiohydrolase (CBH), ß-glucosidase (BG), acid phosphatase (PHOS), N-acetyl-ß-glucosaminidase (NAG) and leucine aminopeptidase (LAP) were assessed fluorometrically using the substrates β-D-cellobioside, β-D-glycopyranoside, phosphate, N-acetyl-β-D-glucosaminide and L-Leucine, respectively. The oxidative enzymes peroxidase (OX1) and phenol oxidase (OX2) were measured colorimetrically with the substrates 3,3’, 5,5’-Tetramethylbenzidine (TMB + H_2_O_2_)^88^ and 2,2’-azino-bis(3-ethylbenzothiazoline-6-sulphonic acid) (ABTS)^89^, respectively. Fluorescence (hydrolytic) and absorbance (oxidative) were measured on a BioTek Synergy HT Multi-detection Microplate Reader at 460 nm and 450 nm, respectively. Final enzyme activity values were calculated as in DeForest^90^ and are reported as μmol substrate h^−1^ g^−1^ dry soil.

To characterize isolate biomass chemistry and melanin content, four replicates of each isolate were grown in potato dextrose broth media to facilitate biomass collection. The liquid media contained the same concentrations of D-glucose, potato infusion extract, MES buffer and ammonium nitrate (C:N = 20) as the substrate media used in the model soil experiments, and the pH was also adjusted to 4.5. Cultures were inoculated into sterilized glass vials containing 100 ml liquid media and shaken continuously at 75 rpm^32^. Cultures were grown at 25°C for between 3-30 days depending on species’ growth rate. Fungal biomass was collected by vacuum filtration and washed three times with FDIW (100 ml/rinse). Biomass was subsequently dried at 70°C and subsampled for biomass chemistry and melanin analyses. The chemical composition of isolate biomass (ground subsamples) was characterized by pyrolysis gas chromatography-mass spectrometry (Py-GC/MS), following methods described previously^91,92^, and the relative proportions of broad chemical compound classes (e.g., polysaccharides, aromatics, lipids, proteins) were determined (additional details below). Biomass melanin concentrations were measured by quantitative colorimetric analysis with Azure A dye^93^. Briefly, dried biomass was incubated for 90 minutes in an Azure A solution (initial absorbance = 0.665 at 610 nm) and subsequently filtered through a 0.45 μm nitrocellulose syringe tip filter. Absorbance of the filtrate was measured with a spectrophotometer at 610 nm and the change in absorbance was compared against a standard curve to calculate biomass melanin content. The standard curve was constructed using known amounts of melanin isolated from *Cenococcum geophilum* biomass, as in^93^.

### Total C, MAOM-C and POM-C analyses

Air-dried soil (5 g) from the long-term incubation study was separated into MAOM (< 53 μm) and POM (> 53 μm) fractions using a standard size-based fractionation protocol^94,95^. Soils were dispersed via shaking with dilute (0.5%) sodium hexametaphosphate for 18 h and were subsequently rinsed through a 53 μm sieve. The MAOM fraction underwent an additional rinse step, followed by centrifugation, to remove DOC that may have been present in the water column. MAOM and POM fractions were oven dried at 105°C and 60°C, respectively. Subsamples of MAOM, POM and whole soil were finely ground and analyzed for total C and N by dry combustion analysis (Perkin-Elmer Series II 2400 Elemental Analyzer; Perkin Elmer Inc.).

### Stable aggregate formation

The potential of each fungal species to generate water-stable aggregates was assessed using a standard wet-sieving procedure (USDA NRCS, 1999). Since the model soil mixture began as a combination of un-aggregated particle separates (sand, silt, clay), any aggregates that formed could be attributed to the growth and activity of the fungal isolates. After harvest of the long-term incubations, soils (5 g) were weighed onto a 250 μm sieve and wetted through capillary action (5 minutes) by positioning the surface of the sieve screen just above water-level within a plastic bin containing ∼1 inch DI water. Thereafter, soils were wet-sieved for three minutes. Deionized water was added to the bin such that the soil on the sieve was completely submerged. The sieve was moved up and down over a vertical distance of 1.5 cm at a rate of 30 oscillations per minute. At the end of the three-minute period, aggregates remaining on the sieve surface were transferred to an aluminum drying pan and oven-dried at 105°C. Water-stable aggregates were calculated as the proportion of the initial 5 g soil that remained on the 250 μm sieve after wet-sieving, minus the amount of remaining soil > 250 μm present in sterile control samples (Sample_(% of soil > 250 μm)_ – Control_(%soil > 250 um)_).

### Chemical and biological stability of SOM

We determined the chemical stability of fungal-derived SOM using a sequential chemical extraction procedure. Each soil sample was subjected to three sequential C-free extractants to isolate discrete pools of SOM that form distinct bonds and/or associations with soil minerals^96,97^. Water was used first to extract readily solubilized compounds, followed by KCl (1 M; pH = 4.8) to isolate easily exchangeable pools, and finally by sodium pyrophosphate (0.1 M; pH 10) to extract compounds bound via stronger electrostatic forces or within organo-metal complexes. For each extraction, 30 ml extraction reagent was added to 1 g soil, vortexed, and shaken for 4 hours (water, KCl) or 16 hours (sodium pyrophosphate) at 200 rpm^96,97^. Samples were centrifuged and the supernatant was filtered using 0.2 μm nylon syringe tip filters. Extracts were analyzed for total organic C (TOC) by high temperature catalytic oxidation followed by non-dispersive infrared detection with a Shimadzu TOC analyzer (TOC-LSH; Shimadzu Corporation, Kyoto, Japan). The percent of initial soil C (pre-extraction) removed by each extractant was calculated for each isolate (full results included in supplementary materials; Fig. S12). The proportion of initial C remaining in the soil pellet (that is, C not removed by the three extractants) was calculated as a metric of SOM chemical stability (% “chemically stable” C). These values are reported as (sample-control) to account for the small amount of C remaining in uninoculated control samples after sequential extraction, likely representing the fraction of substrate C that was directly sorbed to minerals.

We also assessed the biological stability of fungal-derived SOM in a separate 3-month incubation (*sensu* Kallenbach et al.^14^). After soils were harvested from the long-term incubation experiment (described above), 10 g soil subsamples (4 replicates/isolate) were re-inoculated with a mixed microbial community and ^13^C-glucose (to stimulate decomposition), and the relative stability of SOM generated by each isolate was assessed. Since unlabeled (^12^C) glucose was used as a substrate in the original incubation experiment, we applied ^13^C-labeled glucose (25 atom%; 50 μg C g^−1^ soil, as in Kallenbach et al.^14^) in the 3-month biological stability assay, such that we could parse microbial respiration derived from the newly added glucose (^13^CO_2_) versus existing fungal-derived SOM (^12^CO_2_). The microbial inoculum was prepared from soil collected from control plots at the Harvard Forest LTER. Soil was combined with FDIW (1:25 w/v; as in^98^) and the viability of the inoculum was confirmed at pH 4.5 by measuring its respiration for three days following the addition of 120 mg L^−1^ glucose^71^. The microbial inoculum was added in 1 ml aliquots to each 10 g sample, and additional FDIW was added to bring soil moisture back up to 50% field capacity. Soils were incubated in mason jars at 25°C and were hand-mixed once weekly to encourage C mineralization^14^. A water control was established alongside each ^13^C-glucose amended sample to represent background 8^13^C values. After 3 mo. incubation, soils were harvested and analyzed for 8^13^C and total C (%) at the Cornell Stable Isotope Laboratory. A standard isotope mixing model^99^ was used to calculate the relative proportion of isolate-derived SOM that resisted decomposition by the mixed microbial community (% “biologically stable” C). These values are reported as (sample-control) as described above for chemically stable C.

### Chemical composition of fungal-derived SOM

We characterized the chemistry of fungal-derived SOM using Py-GC/MS, following methods described in^91,92^. Subsamples of whole soil and the MAOM fraction were analyzed separately for each isolate. MAOM fraction samples were pyrolyzed at a single temperature (600°C) on a CDS Pyroprobe 5150 pyrolyzer (CDS Analytical Inc., Oxford, PA), following standard protocol, whereas whole soil samples were analyzed using a modified ramped pyrolysis procedure (*sensu*^48^), whereby each sample was pyrolyzed at increasing temperatures (330, 396, 444, 503, 600 and 735°C) in a stepwise fashion. This approach characterizes the composition of organic compounds that require different activation energies for thermal decomposition as a means of assessing the relative thermal stability of SOM^48^. Following pyrolysis (of MAOM or whole soil samples), decomposed products were transferred to a Thermo Trace GC Ultra gas chromatograph (Thermo Fisher Scientific, Austin, TX) and subsequently to a Polaris Q mass spectrometer (Thermo Fisher Scientific). Products were ionized and detected using AMDIS (V. 2.69) and recorded peaks were classified against the National Institute of Standards and Technology (NIST) compound library. Relative percentages of organic matter compounds were summarized within broad compound classes (polysaccharides, lipids, aromatics, phenolics, proteins, N-bearing (non-proteins) or unknown source)^91^.

### Statistical analyses

All statistics were performed in R 4.3.0 unless stated otherwise. Differences in species’ trait values and contributions to SOM functional pools were evaluated using one-way ANOVA and post-hoc Tukey HSD tests (*P* < 0.05). Levene’s test of homogeneity of variances and the Shapiro-Wilk normality test were used to assess homoscedasticity and normality of residuals, respectively. When normality or homoscedasticity assumptions were not met, data were evaluated using non-parametric Kruskal-Wallis and post-hoc Dunn tests, or were square-root or log-transformed when used in other analyses (e.g., regressions).

We used partial least squares regression (PLSR) to evaluate the relative importance of fungal traits associated with the formation of different SOM functional pools. A separate PLSR model was constructed for each SOM pool (response variable), and all measured fungal trait variables were included as predictors. PLSR was conducted in JMP Pro (version 16.1.0) using the NIPALS algorithm (as in^100^). Due to the nature of our dataset, where some of the trait measurements were collected on independent experimental units separate from the long-term incubated samples analyzed for SOM, we used a randomization approach with three iterations to construct our final PLSR model results. We ran three separate PLSR analyses, each run on a separate randomized dataset representing the three possible pairings between each trait measurement replicate (3 replicates per isolate, representing the minimum number of replicates available for certain traits) and each SOM functional pool replicate (3 replicates/isolate) (full model results in Table S3). After model results were collated, we calculated the average X loadings and variable importance in projection (VIP) scores for each trait variable across the three PLSR model iterations. Predictor variables with a VIP score greater than 0.8 were retained in the final models. Thereafter, we fit linear regressions between SOM functional pools and the trait variable with the strongest loading on each pool’s most explanatory PLSR latent factor, and we used ANOVA to assess the significance of these relationships.

Principal components analysis (PCA) was used to summarize fungal trait variables along major axes of variance. PCA was conducted using the ‘rda’ function in the ‘vegan’ package^101^ on z-score standardized values for each trait variable. Principle component axes were retained if eigenvalues were greater than one, and such that cumulative variation explained reached 80%. For ease of interpretation, only the two most explanatory axes (PC1 and PC2), explaining a total of 45% of variation in fungal trait data, are presented and visualized using ordination.

Multivariate ANOVA (MANOVA) was used to test for statistical differences among species and phyla (‘adonis2’ function, Euclidian distance method). Additionally, we examined relationships between PC1 and SOM functional pools by fitting polynomial regression models with the lowest mean square error between each predictor and response variable. When the single degree model (h = 1) exhibited the lowest mean square error value, linear regression was used.

To generate a metric of trait multifunctionality as well as SOM formation and stabilization potential (essentially, SOM multifunctionality) for each isolate, we adapted an approach for calculating “ecosystem multifunctionality”^46,102^. Broadly, multifunctionality is defined as the simultaneous performance or provisioning of multiple functions^46^. We used a modified version of the averaging approach^46^ that uses Hill numbers (q) to produce a single metric of “effective” multifunctionality, which accounts for both the total provisioning of functions as well as the evenness (e.g., relative trait values) of those provisions of function^47^. Trait (or SOM) multifunctionality was calculated using the ‘multifunc’ package (v. 0.9.4)^46,47^ as the product of the effective number of functions performed by each isolate and the arithmetic mean performance of the measured functions. Values for turnover length (originally reported in days) were “reflected” such that the maximum level of each trait variable in our dataset represented the “best” or highest function. We multiplied turnover length values by −1 and then added the un-reflected maximum value such that the lowest level of transformed function was zero^46^. All trait values were then standardized by the maximum observed value to a common scale between 0 to 1 before analysis. Effective multifunctionality was calculated at order q=1 to accommodate information about unequal levels of functioning proportional to the relative functional performance, without making any uncomfortable decisions about upweighting high performing functions^47^. Radar plots were used to visualize trait performance across isolates, with data scaled between 0 to 1 prior to plotting, and differences in effective multifunctionality were evaluated using ANOVA.

Finally, we assessed the effect of isolate identity on SOM and MAOM chemical composition using permutational multivariate analysis of variance (PERMANOVA)^103^ on a Bray-Curtis dissimilarity matrix calculated from the relative abundance values of individual compounds identified by Py-GC/MS. For the whole soil samples that were analyzed with the ramped pyrolysis approach, separate PERMANOVA analyses were run for each thermal fraction. To visualize the effects of species identity on SOM chemistry, we conducted non-metric multidimensional scaling (NMDS) ordination of the Bray-Curtis distance values. Environmental vectors representing broad Py-GC/MS compound classes were fitted to the NMDS using the ‘envifit’ function (‘vegan’ package) and were included in the plot if significant (*P* < 0.05). Lastly, we calculated SOM chemical diversity for each species within the most thermally stable fraction (735°C) of SOM using the Shannon index (individual compound-level dataset).

## Acknowledgements

We thank Jessica Ernakovich, Marco Keiluweit, and Rich Smith for providing feedback on the earlier stages of this manuscript. Thank you also to Mel Knorr for laboratory support, as well as Andrea Jilling and Noah Sokol for feedback on experimental design and analytical approaches. This work was supported by a University of New Hampshire Dissertation Year Fellowship to E.D.W., as well as a research grant from the U.S. Department of Agriculture National Institute of Food and Agriculture through the New Hampshire Agricultural Experiment Station (NHAES; Hatch NH-00701). This is NHAES Scientific Contribution Number 3001.

## Author contributions

E.D.W., A.S.G. and S.D.F. conceived of the study, and K.G. and E.W.M. contributed to early experimental design and methodological development. E.D.W. led laboratory work, sample analysis, data analysis and writing. All authors contributed to data interpretation and editing of the manuscript.

## Supplementary Methods

### Key experimental design decisions and rationale

#### Pilot experiments to reduce contamination risk

A series of pilot experiments were conducted prior to beginning the final experiment described in this manuscript. The primary goals of the pilot experiments were (1) to ensure that all fungal isolates could be grown in the model soil environment with glucose as a primary C source, and (2) to evaluate methodological approaches that reduced the risk of contamination, such that we could ensure single-species axenic conditions in the full experiment. The greatest risks of contamination occurred during respiration measurements, jar flushing (O_2_ replenishment) and substrate amendments. Sterile culturing techniques were used during incubation set-up, and all tools, model soils, incubation containers (jars, specimen cups) and substrate media were autoclaved prior to set-up. All work was conducted within a Laminar flow hood recommended for microbial culturing. Through pilot experiments, we determined that using sterile single-use syringes and needles to make respiration measurements separately for each isolate (different syringe utilized for each grouping of experimental units associated with an isolate on each sampling day) reduced contamination risk. Rather than opening jars to replenish oxygen, jars were flushed using sterile needles fitted with 0.2 μm filters attached to tubes supplying CO_2_-free air. The 0.2 μm filters were labeled by species and were re-utilized only for experimental units of the same isolate. Needles (sterile) were replaced between each instance of flushing. Substrate amendments over the course of the long-term incubation experiment (3-6 months) posed a particularly unique challenge to the maintenance of axenic conditions. We developed an approach utilizing a custom 8-inch stainless steel needle that could be fitted to a glass syringe so that substrate amendments could be made without opening jars. The 8-inch needle was inserted through the rubber ports fitted in the lids of each mason jar, with this particular length being selected such that the needle could reach the bottom of each specimen cup containing model soils. This allowed us to inject the sterilized substrate solution into soils while moving the needle steadily over the vertical length of each soil “core” (50 g cup of soil). The stainless-steel needles and glass syringes were autoclaved (sterilized) prior to substrate amendments.

Based on the results of pilot studies, we decided to establish additional replicates for each isolate to ensure that sufficient replication would remain in case of contamination. A total of eight replicates were established for each isolate. The species that was slowest to establish during initial growth and biomass production, *Panellus stipticus*, was the most challenging taxa to maintain sterile conditions for, with most of the contamination issues occurring during early incubation before *P. stipticus* had established substantial biomass. For this species, we only successfully maintained three experimental units (replicates) under axenic conditions, and therefore were only able to include three replicates in subsequent statistical analyses. For all other isolates, we maintained axenic conditions for at least four replicates, with most species having 5-8 replicates remaining. When > 4 axenic replicates remained, four replicates were selected at random (random number generator) for inclusion in statistical analyses.

#### Standardization approach based on isolate growth dynamics

Various possible approaches to standardize the timing of substrate amendments and incubation length/harvests were considered in the experimental design phase. An obvious alternative would be to standardize by time (e.g., Domeignoz-Horta et al., 2021). In such a scenario, substrate amendments would be made at the same time for all isolates, and experimental units would be incubated for the same amount of time (e.g., experimental units for all isolates harvested at 3 months). Such an approach would provide insight into differences in fungal species’ contributions to SOM formation and stabilization per unit time, and would be especially relevant if the primary research question was focused on the rate of SOM formation by different fungal isolates. However, the primary aim of our experiment was to evaluate relationships between the trait profiles of fungi and their contributions to different pools of SOM, including relatively stable SOM pools. Had all species been incubated for the same amount of time, it is possible that slower growing species may not have utilized all the added substrate-C, such that unprocessed glucose remained in the soil for some isolates, but not others. This would have (1) biased the measurements of species’ contributions to SOM functional pools, especially chemically and biologically stable pools of C, and (2) would have biased our interpretations of trait-SOM relationships. By standardizing by time, it is likely that the traits of fast-growing species with high biomass production would have emerged as important positive predictors of SOM formation (especially stable SOM formation), even if these traits declined in importance over time.

We developed a strategic study design to promote maximum substrate utilization by fungal isolates, such that we could assume a majority of substrate-C had been utilized prior to incubation harvest. We determined the timing of substrate amendments and incubation harvest independently for each isolate. The slowest growing species were incubated for up to ∼6 months, whereas the fastest growing species were harvested at ∼3 months. Our approach was based on monitoring isolate respiration rates, such that each isolate went through the same pattern of growth dynamics before incubation harvest (described in main text). Based on the maximum respiration rates observed across isolates following substrate amendments, we selected a value of ≤ 1 μg CO_2_-C g^−1^ soil hour^−1^ to represent a near-zero respiration rate (see Fig. S3). For all fungal isolates included in this study, peak respiration (after substrate amendment) was followed by declining respiration values, and steady-state growth was determined when isolates exhibited consistently low respiration rates for an extended period of time (≤1 μg CO_2_-C g^−1^ soil hour^−1^). At this point, it was determined that isolates had utilized a majority of the added substrate-C, and this benchmark was used to determine the timing of the second and third substrate amendments as well as final incubation harvest (on an isolate-specific basis; see main text for additional details).

Alternative approaches to assess complete utilization of the added substrate-C could include: (1) direct measurement of glucose in soils, or (2) measurement of total fungal biomass C at the end of the incubation, and calculation of total C utilization (total respired C + fungal biomass C). However, (1) glucose measured in soils at the end of the experiment would not necessarily reflect unutilized substrate-C, as glucose (or other sugars with similar chemical structures) can be exuded by fungi into the soil environment as exudates or metabolites (Hill et al., 2008). Secondly, (2) we were unable to estimate total fungal biomass in model soils at the end of the incubation experiment due to challenges with the chloroform fumigation extraction (CFE) method (discussed in main text). Researchers have encountered similar problems with the CFE method in past studies involving model soils (e.g., Jilling et al., 2021; Kallenbach et al. 2016), perhaps due to the high abundance of “clean” mineral surfaces within model soils compared to natural soils. Our strategic experimental design (based on isolate-specific respiration monitoring) thus provided the most robust approach to estimate (near-)complete substrate utilization by fungal isolates while the experiment was in progress, allowing us to determine appropriate timing for substrate additions and to assume a majority of the added substrate-C had been utilized prior to incubation harvest.

#### Choice of glucose as a primary C source

While multiple C-supplying substrates were considered for use in this experiment, we ultimately decided to include glucose as the primary C source (90% of C; other 10% supplied by potato dextrose infusion). Glucose has been used in numerous studies as a model substrate and has been shown to be rapidly assimilated and metabolized by a wide range of microbial taxa (Hill et al., 2008), including saprotrophic fungi (Rousk & Baath, 2011). A key aspect of our experimental design was the strategic timing of substrate additions and harvests to promote complete utilization of substrate-C.

Compared to other C-supplying substrates, glucose exhibits low levels of discrimination among microbial taxa (Jones et al., 2018). While more complex substrates (e.g., cellulose) were considered, glucose is water-soluble and therefore could be applied more evenly and homogenously throughout the soil as a liquid solution (autoclaved/sterilized). Together, these decisions guided our selection of glucose as a primary C-supplying substrate.

#### Substrate stoichiometry and carbon application rate

The C:N ratio of the substrate media used in this experiment was similar to soil C:N values observed at the site where fungal cultures were isolated, the Harvard Forest (C:N range from ∼16-27 for mineral soils; Compton & Boone; 2000). We chose a C:N ratio of 20:1 specifically, as it was expected to alleviate N limitation for fungal isolates (Manzoni et al., 2012; Sinsabaugh et al., 2016), whose average biomass C:N ratio was ∼10 (Morrison et al., 2022). A total of three substrate amendments (1 ml each; 5.6 mg C g^−1^ soil) were applied over the course of the incubation. Each experimental unit contained 50 g soil, and thus 240 mg C were added per substrate amendment. The rate of C application was selected to (1) minimize the total number of substrate additions required (lowering contamination risk) and (2) prevent osmotic shock to fungal isolates (a concern with highly concentrated glucose solutions). At the end of the experiment, a total of 840 mg C had been added to each experimental unit. We selected this amount of C as a sufficient stopping point for our experiment, because it was expected to increase soil C concentrations into a range comparable to the C content of agricultural mineral soils (0.8-1.0% C). While this range is low compared to more C-rich forest or grassland soils, it is still a realistic range of total C values observed among natural field soils, and it allowed us to end the incubations after ∼3-6 months (depending on isolate). As discussed in the main text, a central constraint of our study was the need to maintain axenic conditions of fungal cultures. Because the risk of contamination increases with duration of incubation and number of substrate amendments, we decided that three substrate additions (840 mg C) would be sufficient to observe and compare differences in SOM formation across fungal isolates.

#### Multiple approaches to quantify isolate CUE

Given that carbon use efficiency (CUE) measurements can vary based on isolate growth stage and resource availability (e.g., C uptake from glucose vs. biomass recycling), we decided to include three separate metrics of CUE in our analysis of trait-SOM relationships. The three metrics were as follows: CUE measured during log-phase growth in (1) liquid culture and (2) model soils, and (3) CUE measured at stationary growth in model soils. Liquid culture values, previously published by Morrison et al. (2022), were collected from short-term incubations (∼1 week to 1 month, depending on isolate) where respiration and biomass measurements were made at regular intervals. These values are included as a representation of fungal isolates’ maximum CUE under optimal growth conditions. Values ranged from 0.33 ± 0.03 to 0.72 ± 0.08, consistent with previous studies of pure microbial cultures grown on non-limiting supplies of glucose (0.4-0.88), which can approach theoretical maxima (Geyer et al., 2016). The CUEs of fungal isolates in model soils (assessed during log-phase growth) were measured in short-term incubations mirroring the liquid culture approach. Soil CUE (log-phase) values were lower on average than liquid culture measurements, ranging from 0.21 ± 0.03 to 0.60 ± 0.02. These values are more similar to CUE values observed for soil microbial communities (0.24-0.77; reviewed in Geyer et al., 2016), which tend to be lower than maximal values observed in liquid culture, in part because soil properties can limit substrate availability and C uptake rates (e.g., via temporary mineral sorption or occlusion within pore spaces). Both the liquid culture and soil CUE measurements made during log-phase growth are expected to capture CUE before significant biomass turnover begins (Geyer et al., 2019), which is known to influence CUE (Geyer et al., 2016). The third metric of CUE, assessed in model soils during stationary growth, integrates this effect of biomass turnover. This measure of CUE (measured via ^18^O-water incorporation into DNA) captures both the efficiency of substrate retention as fungal biomass, as well as the efficiency of biomass recycling across generations of fungal cells (*sensu* Geyer et al., 2016). Interestingly, soil CUE at stationary growth ranged from 0.46 ± 0.04 to 0.96 ± 0.01, making it the highest of the three CUE measurements (on average). While past studies have shown that biomass turnover and substrate recycling can reduce CUE values for mixed soil microbial communities, especially if microbial necromass becomes incorporated into temporarily inaccessible SOM pools (Geyer et al., 2016), relatively high CUE values observed in our experiment for some isolates could be indicative of more “self-sustaining” fungal populations that efficiently recycle (in this case, their own) biomass C. We may have been more likely to observe this pattern in our experiment (compared to natural field soils) because of the high levels of biomass production per unit area (visible to the naked eye and under a microscope), which likely represented a large pool of substrate available for fungal recycling.

Because each individual metric of CUE is unlikely to represent the long-term CUE dynamics of fungal isolates on its own, we calculated an average value that integrates the three metrics to better approximate mean CUE of fungal isolates over the course of the long-term incubations. These values are integrated into certain analyses presented in the main text (e.g., Fig. 1; Fig. 4). In addition, we presented each of the three CUE metrics separately in analyses examining relationships between fungal traits and SOM formation (e.g., PLSR, Fig. 4) such that readers could assess these relationships individually for each CUE measure x SOM pool combination.

### Modified phenol-chloroform DNA extraction

Anticipating potential challenges with extracting DNA from model soils with a high abundance of “clean” mineral surfaces, we used a modified phenol-chloroform DNA extraction procedure recommended for high clay and Fe-oxide rich soils (Griffiths et al., 2000; Shi et al., 2015) and adapted from previous studies on model soils (e.g., mineral packets; Whitman et al., 2018). All solutions used in this procedure were nuclease-free. Briefly, soils (∼0.3 g) received 1.168 ml urea extraction buffer (4.67 ml g^−1^) and 82.5 μl lysis buffer (4M guanidine isothiocyanate, 1% sarkosyl, 1M NaPO_4_, 10% bovine serum albumin, 10 μl L^−1^ 2-(β)-mercaptoethanol) at pH 8, and lysing matrix E beads (MP Biomedicals, Santa Ana, CA). Tubes were shaken on a Biotech Mini Bead Beater for 2.5 minutes, and subsequently centrifuged. The supernatant was transferred to a new tube and combined with 325 μl detergent solution (5% sarkosyl, 5% CTAB), 168 μl 5M potassium acetate, and an equal volume of 24:1 chloroform/isoamyl alcohol. Samples were centrifuged at 10 K x *g* for 10 minutes, and the supernatant transferred to a new tube and combined with 0.6 volumes of isopropanol and 2 μl GlycoBlue co-precipitant (Invitrogen). Samples were incubated for 1.5 hours at room temperature and subsequently centrifuged for 30 minutes at 16 K x *g*. The resulting supernatant was removed from the tube, leaving only the DNA pellet, which was washed with 1 ml 70% ice-cold ethanol (molecular grade). Samples were centrifuged for 10 minutes at 16 K x *g*, ethanol was removed, and pellets were air-dried and resuspended in 90 μl Tris EDTA buffer. DNA solutions were frozen overnight and purified by incubation with 7.5M LiCl (30 minutes at 4°C) the following day. After incubation, samples were centrifuged, supernatant removed, and combined with 80 μl isopropanol. Samples were incubated for 30 minutes at −20°C, centrifuged, supernatant removed, and the DNA pellet washed with 70% ice-cold ethanol. DNA pellets were dried and then dissolved in 20 μl PCR-grade water. Lastly, DNA was quantified using a Qubit 3.0 Fluorometer (Life Technologies, Grand Island, NY) following the manufacturer’s protocol.

## Supplementary tables and figures

**Table S1.** Average trait values (standard error in parentheses) observed at the phylum-level for Basidiomycota, Ascomycota, and Mucoromycotina. One-way ANOVA (parametric) or Kruskal-Wallis (non-parametric) results are presented for each trait and associated post-hoc comparisons are presented when ANOVA or Kruskal-Wallis results were significant (Tukey HSD for parametric data; Dunn test for non-parametric data). Optimum growth rate (GR) and CUE were conducted in liquid culture during log-phase growth. Soil growth rate and CUE measurements were conducted during log-phase growth unless specified at stationary phase growth. Units are as follows for each trait measurement: growth rate (μg biomass-C produced biomass-C^−1^ hr^−1^), turnover (days), hyphal length (m hyphae g^−1^ dry soil), hyphal surface area (m^2^ g^−1^ dry soil), hyphal diameter (μm), melanin (mg g^−1^ biomass), proteins and other compounds measured by Py-GC/MS (% relative abundance in biomass), potential enzyme activities (μmol substrate h^−1^ g^−1^ dry soil).

|  | *Phyla* |  |  | *ANOVA/*<br>*Kruskal* | *Post-hoc Tukey HSD/Dunn* |  |  |
| --- | --- | --- | --- | --- | --- | --- | --- |
| *Variable* | **Basidiomycota**<br>**(n=8)** | **Ascomycota**<br>**(n=20)** | **Mucoromycotina**<br>**(n=8)** | **(P-values)** | **Basidio-**<br>**Asco** | **Asco-**<br>**Mucor** | **Mucor-**<br>**Basidio** |
| Optimum GR | 0.006 (0.001) | 0.015 (0.001) | 0.080 (0.008) | <0.001*** | 0.007** | 0.002** | <0.001*** |
| Soil GR | 0.004 (0.0003) | 0.023 (0.004) | 0.042 (0.009) | <0.001*** | 0.003** | 0.033* | <0.001*** |
| Optimum CUE | 0.677 (0.04) | 0.568 (0.04) | 0.461 (0.03) | 0.011* | 0.117 | 0.011* | 0.002* |
| Soil CUE | 0.419 (0.08) | 0.491 (0.03) | 0.304 (0.047) | 0.034* | 0.537 | 0.027* | 0.345 |
| Soil CUE (stationary) | 0.726 (0.09) | 0.781 (0.04) | 0.495 (0.03) | 0.019* | 0.364 | 0.003* | 0.021* |
| Turnover | 1.408 (0.83) | 0.900 (0.23) | 1.433 (0.17) | 0.246 | - | - | - |
| BG + CBH | 964.1 (343.1) | 5271 (581.4) | 842.2 (304.0) | <0.001*** | <0.001*** | <0.001*** | 0.399 |
| PHOS | 436.1 (124.2) | 1553 (219.7) | 1889 (233.3) | 0.012* | 0.006** | 0.197 | 0.003** |
| NAG | 1580 (264.1) | 1959 (412.4) | 894.5 (368.9) | 0.292 | - | - | - |
| LAP | 0.00 (0.00) | 909.7 (615.6) | 0.00 (0.00) | 0.097 | - | - | - |
| OX | 43.43 (13.4) | 10.09 (1.15) | 6.536 (1.84) | 0.002** | 0.003** | 0.081 | <0.001*** |
| Hyphal SA | 76.36 (23.1) | 133.6 (22.0) | 784.2 (50.0) | <0.001*** | 0.137 | <0.001*** | <0.001*** |
| Hyphal length | 17.31 (6.22) | 12.90 (1.55) | 17.50 (1.83) | 0.413 | - | - | - |
| Hyphal diam. | 1.535 (0.14) | 2.634 (0.25) | 15.69 (2.24) | <0.001*** | 0.014* | 0.002* | <0.001*** |
| Melanin | 0.038 (0.001) | 0.036 (0.005) | 0.008 (0.003) | <0.001*** | 0.214 | <0.001*** | <0.001*** |
| Phenols | 5.481 (0.399) | 7.108 (0.413) | 10.78 (1.069) | <0.001*** | 0.159 | <0.001*** | <0.001*** |
| Proteins | 6.809 (0.316) | 10.71 (0.709) | 11.80 (0.973) | 0.005** | 0.002** | 0.241 | 0.002** |
| N-bearing | 8.937 (0.831) | 11.06 (0.812) | 14.23 (0.111) | 0.022* | 0.087 | 0.028* | 0.003** |
| Polysacc. | 32.07 (1.961) | 36.81 (1.123) | 29.28 (2.105) | 0.020* | 0.044* | 0.005** | 0.222 |
| Lipids | 35.62 (2.723) | 17.99 (1.621) | 16.56 (6.077) | 0.003** | <0.001*** | 0.486 | 0.004* |
| Aromatics | 2.246 (0.281) | 3.043 (0.236) | 2.926 (0.296) | 0.158 | - | - | - |

**Table S2.** Average contributions to SOM pools (standard error in parentheses) observed at the phylum-level for Basidiomycota, Ascomycota, and Mucoromycotina. *P*-values from one-way ANOVA (parametric) or Kruskal-Wallis (non-parametric) results are presented for each SOM pool and associated post-hoc comparisons are presented when ANOVA or Kruskal-Wallis results were significant (Tukey HSD for parametric data; Dunn test for non-parametric data).

|  | *Phyla* |  |  | *ANOVA/*<br>*Kruskal* | *Post-hoc Tukey HSD/Dunn* |  |  |
| --- | --- | --- | --- | --- | --- | --- | --- |
| *Variable* | **Basidiomycota**<br>**(n=8)** | **Ascomycota**<br>**(n=20)** | **Mucoromycotina**<br>**(n=8)** | **(P-values)** | **Basidio-**<br>**Asco** | **Asco-**<br>**Mucor** | **Mucor-**<br>**Basidio** |
| Total C (%) | 0.730 (0.03) | 0.695 (0.03) | 0.641 (0.03) | 0.301 | - | - | - |
| MAOM-C (%) | 1.65 (0.16) | 1.47 (0.07) | 1.26 (0.06) | 0.045* | 0.425 | 0.278 | 0.041* |
| POM-C (%) | 0.08 (0.006) | 0.285 (0.02) | 0.238 (0.02) | <0.001*** | <0.001*** | 0.416 | 0.004** |
| Water-stable<br>aggregates (%) | 80.21 (1.60) | 83.13 (1.11) | 61.6 (1.20) | <0.001*** | 0.133 | <0.001*** | 0.003** |
| Chemically<br>stable C (%) | 22.59 (5.46) | 54.38 (2.04) | 46.45 (2.61) | <0.001*** | <0.001*** | 0.044* | 0.032* |
| Biologically<br>stable C (%) | 55.25 (4.17) | 69.68 (2.61) | 71.69 (1.79) | 0.012* | 0.004** | 0.303 | 0.003* |

**Table S3.** Full PLSR model outputs for three separate iterations of PLSR on the randomized dataset (three randomizations). Three iterations of PLSR were conducted for each SOM functional pool. The five variables with the highest loadings on PLSR factor 1 and factor 2 are presented for each model, with the X loading value and VIP scores presented in parentheses. Average values were calculated across the three randomizations for the main text (Fig. 4).

| **Randomization # →** | **Randomization 1** |  | **Randomization 2** |  | **Randomization 3** |  |
| --- | --- | --- | --- | --- | --- | --- |
| **Response (Y)** | **Model info** | **Variable (loading/VIP)** | **Model info** | **Variable (loading/VIP)** | **Model info** | **Variable (loading/VIP)** |
| **Total C** | Two factor model<br>(cumulative R<sup>2</sup>Y = 69.2%):<br>-F1: 56.8%<br>-F2: 12.4% | <u>Factor 1 (56.8%):</u><br>CUE soil, <sup>18</sup>O (0.40/1.80)<br>Growth rate, opti. (-0.31/1.08)<br>Biomass polysacc (0.29/1.36)<br>Growth rate, soil (-0.29/1.19)<br>BG (0.28/1.06)<br><u>Factor 2 (12.4%):</u><br>Hyphal SA (0.37/0.89)<br>NAG (-0.35/1.34)<br>Biomass phenol (0.34/0.80)<br>Hyphal length (0.30/1.73)<br>Melanin (-0.30/0.76) | Two factor model<br>(cumulative R<sup>2</sup>Y = 66.9%):<br>-F1: 47.7%<br>-F2: 19.2% | <u>Factor 1 (47.7%):</u><br>CUE soil, <sup>18</sup>O (0.39/1.88)<br>Growth rate, opti. (-0.32/1.13)<br>Growth rate, soil (-0.29/1.17)<br>Hyphal SA (-0.29/1.04)<br>BG (0.28/1.05)<br><u>Factor 2 (19.2%):</u><br>NAG (-0.40/1.48)<br>Phenol (0.32/1.02)<br>Hyphal SA (0.31/1.03)<br>Melanin (-0.30/0.81)<br>PHOS (0.29/0.81) | Two factor model<br>(cumulative R<sup>2</sup>Y = 69.3%):<br>-F1: 47.7%<br>-F2: 21.6% | <u>Factor 1 (47.7%):</u><br>CUE soil, <sup>18</sup>O (0.38/1.69)<br>Growth rate, opti. (-0.33/1.16)<br>Hyphal SA (-0.30/1.05)<br>Growth rate, soil (-0.29/1.19)<br>BG (0.28/1.01)<br><u>Factor 2 (21.6%):</u><br>NAG (-0.42/1.77)<br>Phenol (0.36/1.13)<br>Melanin (-0.34/0.95)<br>Hyphal SA (0.29/1.06)<br>Lipid (-0.29/0.38) |
| **POM-C** | Two factor model<br>(cumulative R<sup>2</sup>Y = 70.1%):<br>-F1: 65.3%<br>-F2: 4.9% | <u>Factor 1 (65.3%):</u><br>Biomass lipid (-0.37/1.56)<br>CBH (0.35/2.01)<br>ABTS (-0.35/1.48)<br>PHOS (0.30/1.47)<br>Biomass polysacc (0.29/1.66)<br><u>Factor 2 (4.9%):</u><br>Biomass phenol (-0.32/0.57)<br>BG (0.30/1.60)<br>Biomass protein (-0.30/0.88)<br>CBH (0.29/2.01)<br>CUE soil, <sup>18</sup>O (0.29/0.73) | Two factor model<br>(cumulative R<sup>2</sup>Y = 68.8%):<br>-F1: 64.3%<br>-F2: 4.5% | <u>Factor 1 (64.3%):</u><br>Biomass lipid (-0.37/1.58)<br>CBH (0.36/2.08)<br>ABTS (-0.34/1.49)<br>PHOS (0.31/1.48)<br>BG (0.30/1.70)<br><u>Factor 2 (4.5%):</u><br>Biomass protein (-0.33/0.96)<br>Biomass N-bearing (-0.32/0.53)<br>Biomass phenol (-0.31/0.63)<br>CUE soil, <sup>18</sup>O (0.31/0.77)<br>CBH (0.30/2.08) | Two factor model<br>(cumulative R<sup>2</sup>Y = 69.8%):<br>-F1: 64.8%<br>-F2: 5.0% | <u>Factor 1 (64.8%):</u><br>CBH (0.37/2.01)<br>Biomass lipid (-0.37/1.57)<br>ABTS (-0.34/1.41)<br>BG (0.32/1.60)<br>PHOS (0.30/1.55)<br><u>Factor 2 (5.0%):</u><br>Biomass protein (-0.39/0.92)<br>Biomass phenol (-0.32/0.50)<br>Biomass N-bearing (-0.31/0.54)<br>CBH (0.31/2.01)<br>CUE soil, <sup>18</sup>O (0.29/0.86) |
| **MAOM-C** | Two factor model<br>(cumulative R<sup>2</sup>Y = 90.4%):<br>-F1: 74.4%<br>-F2: 16.0% | <u>Factor 1 (74.4%):</u><br>CUE soil, <sup>18</sup>O (0.37/1.50)<br>Growth rate, opti. (-0.30/1.06)<br>CUE optimum (0.30/1.40)<br>Biomass polysacc (0.29/1.27)<br>Biomass N-bear (-0.28/1.05)<br><u>Factor 2 (16.0%):</u><br>Melanin (-0.37/0.65)<br>NAG (-0.35/1.35)<br>Hyphal SA (0.34/0.77)<br>Biomass phenol (0.31/0.81)<br>Hyphal length (0.27/1.95) | Two factor model<br>(cumulative R<sup>2</sup>Y = 90.8%):<br>-F1: 62.4%<br>-F2: 22.8%<br>-F3: 5.5% | <u>Factor 1 (62.4%):</u><br>CUE soil, <sup>18</sup>O (0.35/1.63)<br>Growth rate, opti. (-0.33/1.12)<br>CUE optimum (0.30/1.43)<br>Biomass polysacc (0.29/1.23)<br>Biomass N-bear (-0.29/1.16)<br><u>Factor 2 (22.8%):</u><br>Melanin (-0.41/0.89)<br>NAG (-0.40/1.44)<br>Hyphal SA (0.29/0.96)<br>Biomass phenol (0.30/0.94)<br>BG (-0.28/0.72) | Two factor model<br>(cumulative R<sup>2</sup>Y = 89.7%):<br>-F1: 63.3%<br>-F2: 20.5%<br>-F3: 5.9% | <u>Factor 1 (63.3%):</u><br>CUE soil, <sup>18</sup>O (0.35/1.47)<br>Growth rate, opti. (-0.32/1.12)<br>CUE optimum (0.30/1.33)<br>Biomass N-bear (-0.30/1.26)<br>Biomass polysacc (0.28/1.16)<br><u>Factor 2 (20.5%):</u><br>Melanin (-0.40/0.88)<br>NAG (-0.38/1.57)<br>Hyphal SA (0.29/0.94)<br>Biomass phenol (0.29/0.92)<br>BG (-0.28/0.75) |
| **Chemically stable C** | Two factor model<br>(cumulative $R^2Y = 93.6\%$):<br>-F1: 78.0%<br>-F2: 8.9%<br>-F3: 6.5% | <u>Factor 1 (78.0%)</u><br>Biomass lipid (-0.38/1.67)<br>Biomass protein (0.34/1.53)<br>ABTS (-0.32/1.47)<br>TMB (-0.31/1.16)<br>Biomass phenol (0.28/1.03)<br>Growth rate, soil (0.26/1.11) | Two factor model<br>(cumulative $R^2Y = 92.5\%$):<br>-F1: 75.1%<br>-F2: 9.7%<br>-F3: 7.7% | <u>Factor 1 (75.1%)</u><br>Biomass lipid (-0.38/1.54)<br>Biomass protein (0.35/1.44)<br>ABTS (-0.30/1.17)<br>TMB (-0.30/1.04)<br>Biomass phenol (0.27/1.04)<br>Growth rate, soil (0.27/1.14) | Two factor model<br>(cumulative $R^2Y = 88.3\%$):<br>-F1: 79.6%<br>-F2: 8.7% | <u>Factor 1 (79.6%)</u><br>Biomass lipid (-0.38/1.58)<br>Biomass protein (0.35/1.57)<br>ABTS (-0.32/1.66)<br>TMB (-0.31/1.18)<br>PHOS (0.29/1.37)<br>Biomass phenol (0.28/1.10) |
| | | <u>Factor 2 (8.9%)</u><br>BG (0.36/1.07)<br>Growth rate, opti. (-0.36/0.60)<br>CUE soil, $^{18}O$ (0.35/0.51)<br>Biomass polysacc (0.33/1.09)<br>Hyphal SA (-0.33/0.64) | | <u>Factor 2 (9.7%)</u><br>BG (0.35/1.11)<br>Growth rate, opti. (-0.35/0.59)<br>CUE soil, $^{18}O$ (0.33/0.54)<br>Biomass polysacc (0.28/0.96)<br>Hyphal SA (-0.38/0.54) | | <u>Factor 2 (8.7%)</u><br>BG (0.35/1.00)<br>Growth rate, opti. (-0.35/0.55)<br>CUE soil, $^{18}O$ (0.33/0.41)<br>Biomass polysacc (0.29/0.80)<br>Hyphal SA (-0.37/0.55) |
| **Biologically stable C** | Two factor model<br>(cumulative $R^2Y = 68.6\%$):<br>-F1: 47.5%<br>-F2: 21.1% | <u>Factor 1 (47.5%)</u><br>Biomass protein (0.35/1.73)<br>Biomass phenol (0.33/1.17)<br>Growth rate, soil (0.31/1.26)<br>TMB (-0.28/1.18)<br>Biomass N-bear. (0.27/1.14) | Two factor model<br>(cumulative $R^2Y = 63.8\%$):<br>-F1: 40.1%<br>-F2: 12.0%<br>-F3: 11.8% | <u>Factor 1 (40.1%)</u><br>Biomass protein (0.37/1.42)<br>Biomass phenol (0.33/1.03)<br>Growth rate, soil (0.31/1.20)<br>Biomass lipid (-0.31/1.15)<br>Biomass N-bear. (0.27/1.09) | Two factor model<br>(cumulative $R^2Y = 64.4\%$):<br>-F1: 43.5%<br>-F2: 21.0% | <u>Factor 1 (43.5%)</u><br>Biomass protein (0.35/1.73)<br>Biomass phenol (0.33/1.16)<br>Biomass lipid (-0.32/1.39)<br>Growth rate, soil (0.29/1.35)<br>Biomass N-bear. (0.28/1.08) |
| | | <u>Factor 2 (21.1%)</u><br>BG (0.39/0.58)<br>Hyphal SA (-0.37/0.98)<br>Growth rate, opti.(-0.31, 0.96)<br>CUE soil, $^{18}O$ (0.29/0.76)<br>CUE soil (0.27/1.05) | | <u>Factor 2 (12.0%)</u><br>BG (0.42/1.00)<br>Hyphal SA (-0.39/0.91)<br>Growth rate, opti. (-0.32, 0.93)<br>Hyphal length (-0.32/1.37)<br>CUE soil, $^{18}O$ (0.28/1.14) | | <u>Factor 2 (21.0%)</u><br>BG (0.41/0.90)<br>Growth rate, opti. (-0.30, 0.93)<br>CUE soil, $^{18}O$ (0.35/0.80)<br>CUE soil (0.32/1.23)<br>Biomass polysacc (0.29/0.90) |
| **Water-stable aggregates** | Two factor model<br>(cumulative $R^2Y = 89.6\%$):<br>-F1: 75.9%<br>-F2: 13.6% | <u>Factor 1 (75.9%)</u><br>Growth rate, opti. (-0.36/1.72)<br>Hyphal SA (-0.34/1.59)<br>CUE soil, $^{18}O$ (0.33/1.34)<br>Biomass phenol (-0.29/1.13)<br>BG (0.27/1.22) | Two factor model<br>(cumulative $R^2Y = 89.0\%$):<br>-F1: 78.4%<br>-F2: 10.6% | <u>Factor 1 (75.9%)</u><br>Growth rate, opti. (-0.37/1.73)<br>Hyphal SA (-0.37/1.81)<br>CUE soil, $^{18}O$ (0.32/1.36)<br>Biomass phenol (-0.29/1.22)<br>BG (0.29/1.28) | Two factor model<br>(cumulative $R^2Y = 87.7\%$):<br>-F1: 75.2%<br>-F2: 12.5% | <u>Factor 1 (75.9%)</u><br>Growth rate, opti. (-0.37/1.69)<br>Hyphal SA (-0.37/1.83)<br>CUE soil, $^{18}O$ (0.32/1.40)<br>Biomass phenol (-0.29/1.19)<br>BG (0.28/1.23) |
|  |  | <u>Factor 2 (13.6%)</u><br>CUE, soil (0.38/0.94)<br>PHOS (0.32/0.45)<br>LAP (0.31/0.52)<br>ABTS (-0.30/0.64)<br>Growth rate, opti. (0.29/0.99) |  | <u>Factor 2 (13.6%)</u><br>CUE, soil (0.30/0.94)<br>PHOS (0.33/0.49)<br>LAP (0.33/0.64)<br>Growth rate, soil (0.31/1.46)<br>Biomass protein (0.30/0.68) |  | <u>Factor 2 (13.6%)</u><br>CUE, soil (0.32/1.55)<br>PHOS (0.34/0.52)<br>LAP (0.34/0.69)<br>Growth rate, soil (0.32/0.94)<br>Biomass protein (0.29/0.72) |

**Figure S1.**
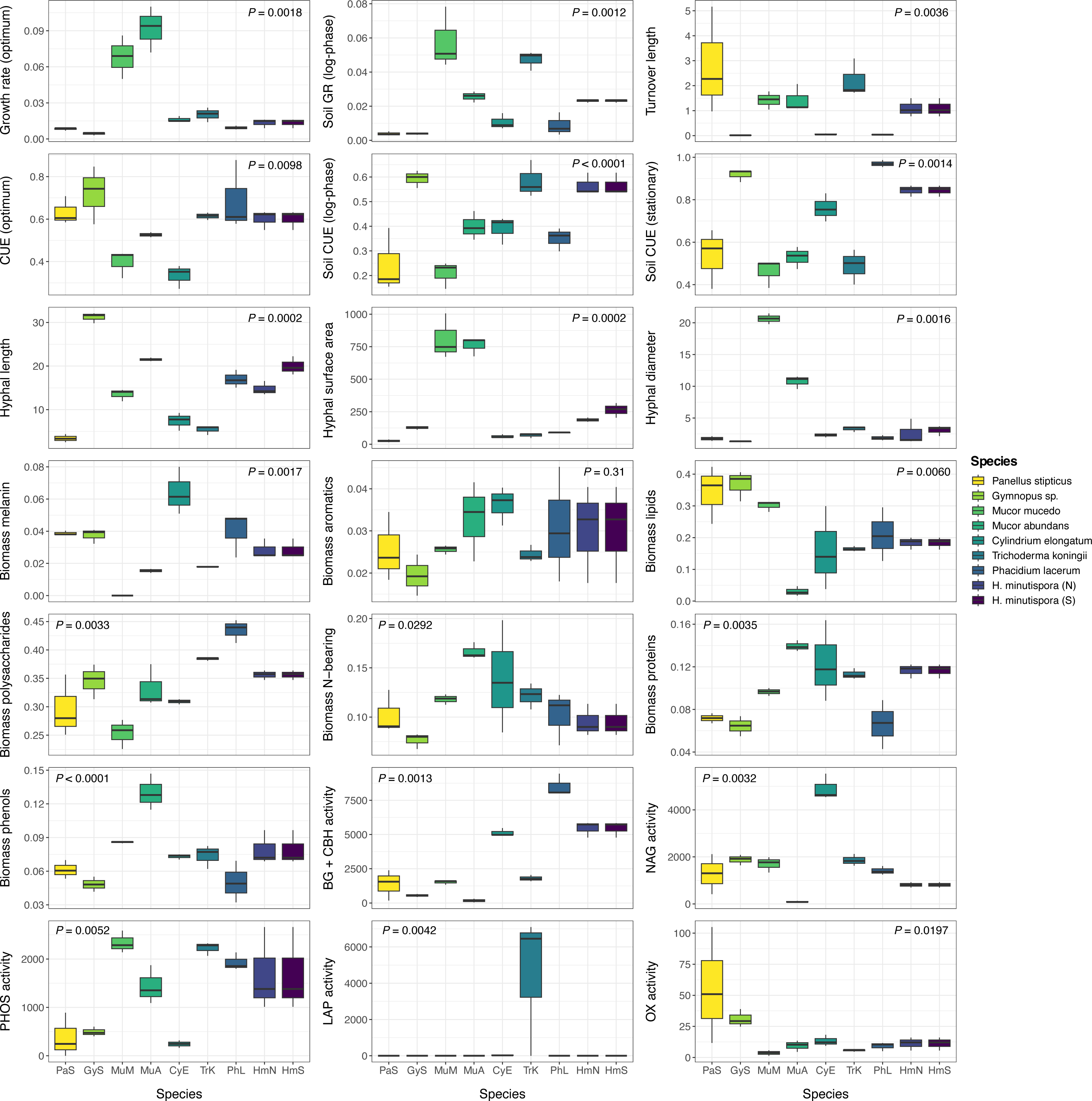
Boxplots showing all measured traits and their relative expression across fungal species. Boxplots represent 25^th^ and 75^th^ percentile, median and outlying points. Results (p-values) are presented for one-way ANOVA or Kruskal-Wallis analyses on the effect of species on each trait measurement (n=4 replicates per species). Optimum growth rate (GR) and CUE were conducted in liquid culture during log-phase growth. Soil growth rate and CUE measurements were conducted during either log-phase or stationary phase growth, as specified in the axis titles. Units are as follows for each trait measurement: growth rate (μg biomass-C produced biomass-C^−1^ hr^−1^), turnover (days), hyphal length (m hyphae g^−1^ dry soil), hyphal surface area (m^2^ g^−1^ dry soil), hyphal diameter (μm), melanin (mg g^−1^ biomass), proteins and other compounds measured by Py-GC/MS (% relative abundance in biomass), potential enzyme activities (μmol substrate h^−1^ g^−1^ dry soil). For *H. minutispora*, samples were separately categorized as sporulating (S) or non-sporulating (N).

**Figure S2.**
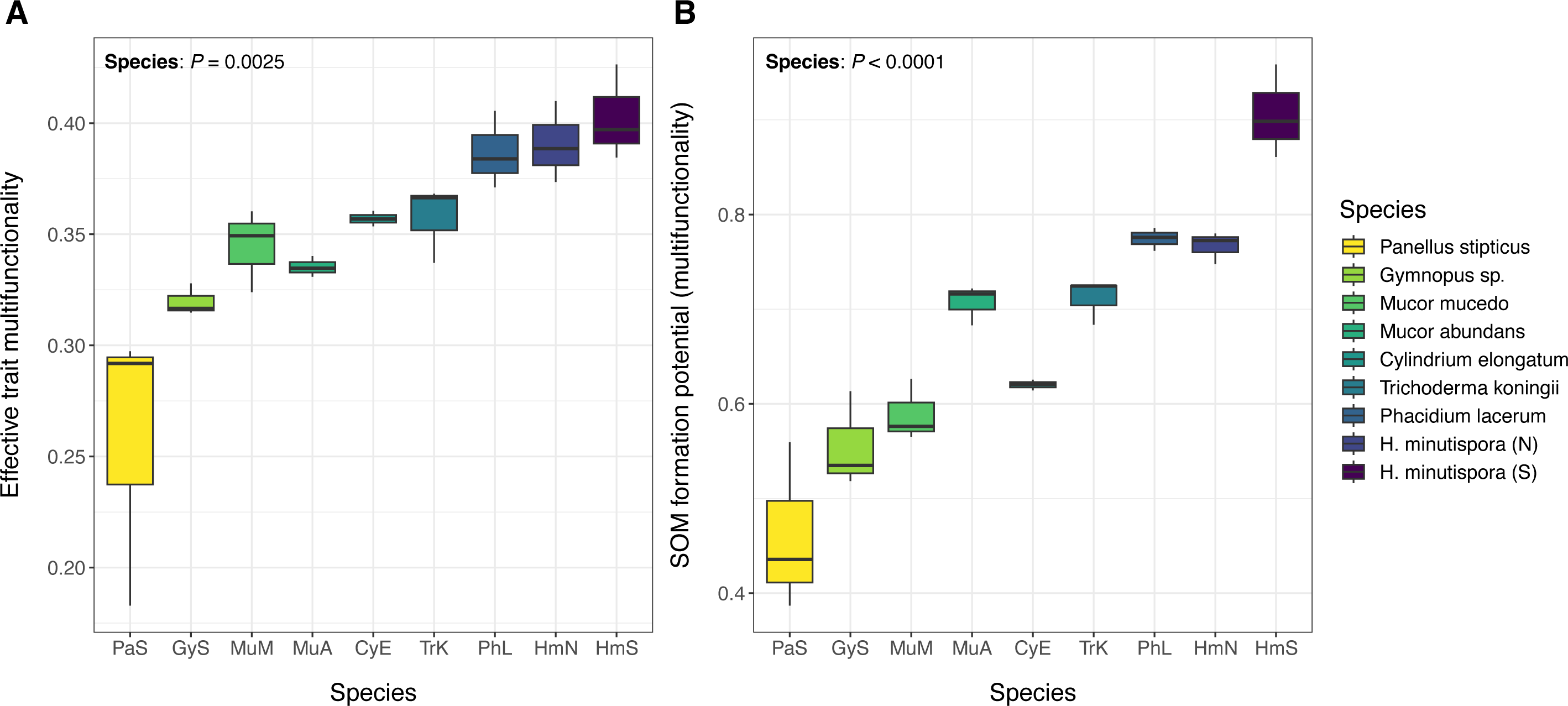
Boxplots of effective trait multifunctionality (A) and SOM formation potential (B) across fungal species. Boxplots represent 25^th^ and 75^th^ percentile, median and outlying points.

**Figure S3.**
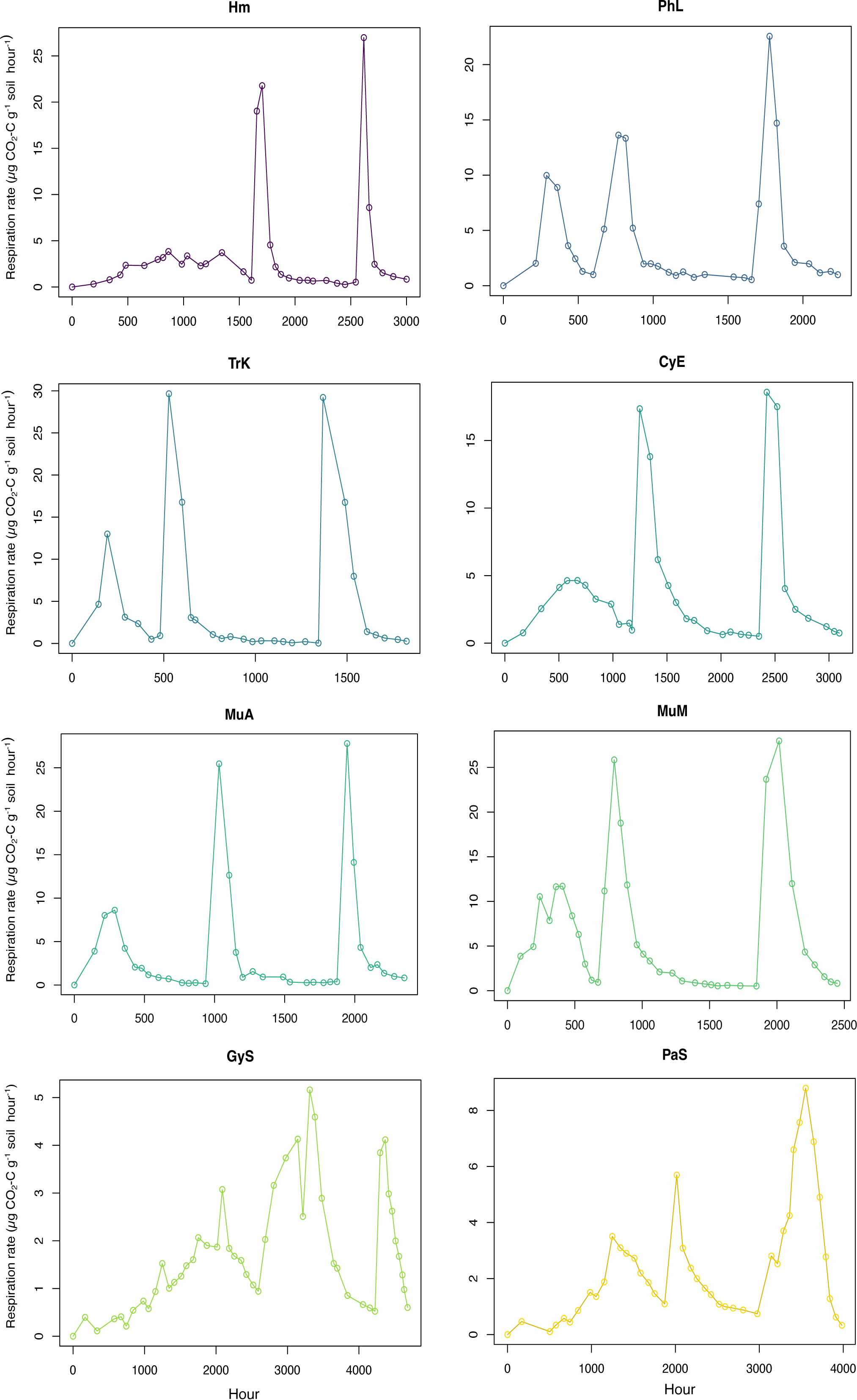
Average respiration rates observed for each fungal species over the course of the long-term incubation. Respiration rates were monitored to determine the timing of the second and third substrate additions and final incubation harvests on an isolate-specific basis. The second substrate amendment was added once an isolate’s respiration rate dropped to ≤ 1 μg CO_2_-C g^−1^ soil hour^−1^. The third substrate amendment was added after isolates had undergone a period of stationary growth for ∼20 days (500 hours), which was determined to begin when each isolate’s respiration rate reached ≤ 1 μg CO_2_-C g^−1^ soil hour^−1^. This period of stationary growth was intentionally induced to promote fungal biomass recycling. After the third substrate addition, isolates underwent an exponential growth phase, followed by a period of declining growth. Final incubation harvests were made once an isolate’s respiration rate had once again declined to < 1 μg CO_2_-C g^−1^ soil hour^−1^.

**Figure S4.**
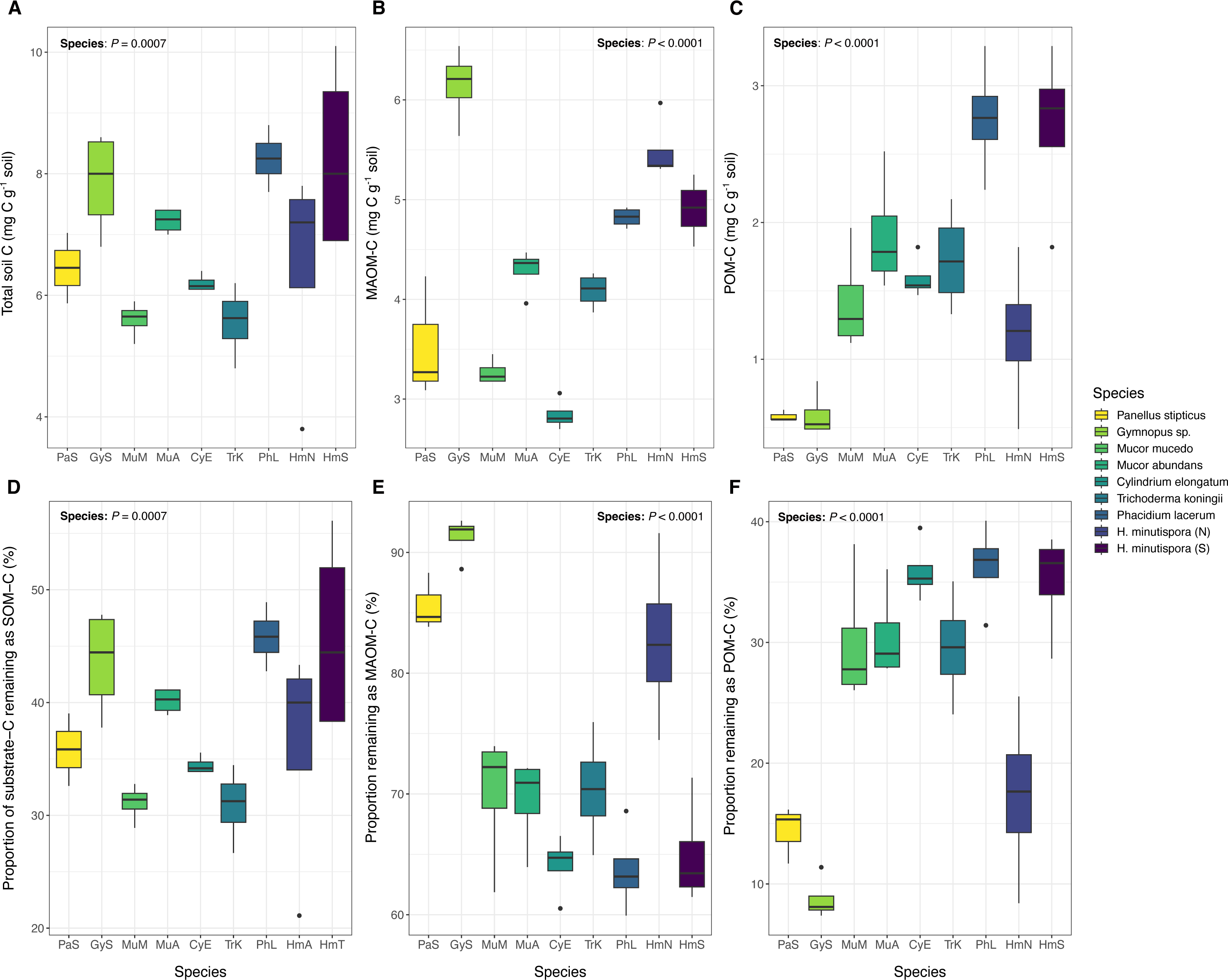
Total soil C concentrations (mg C g^−1^ soil) of bulk soils at incubation harvest (A), and C concentrations within MAOM and POM fractions (mg C g^−1^ soil) (B-C). Proportion (%) of total substrate C added over the course of the long-term experiment that remained as SOM-C (panel D); (E) proportion of remaining soil C that was present in the MAOM fraction, compared with the POM fraction (F). Colors represent individual fungal species. Boxplots represent 25^th^ and 75^th^ percentile, median and outlying points.

**Figure S5.**
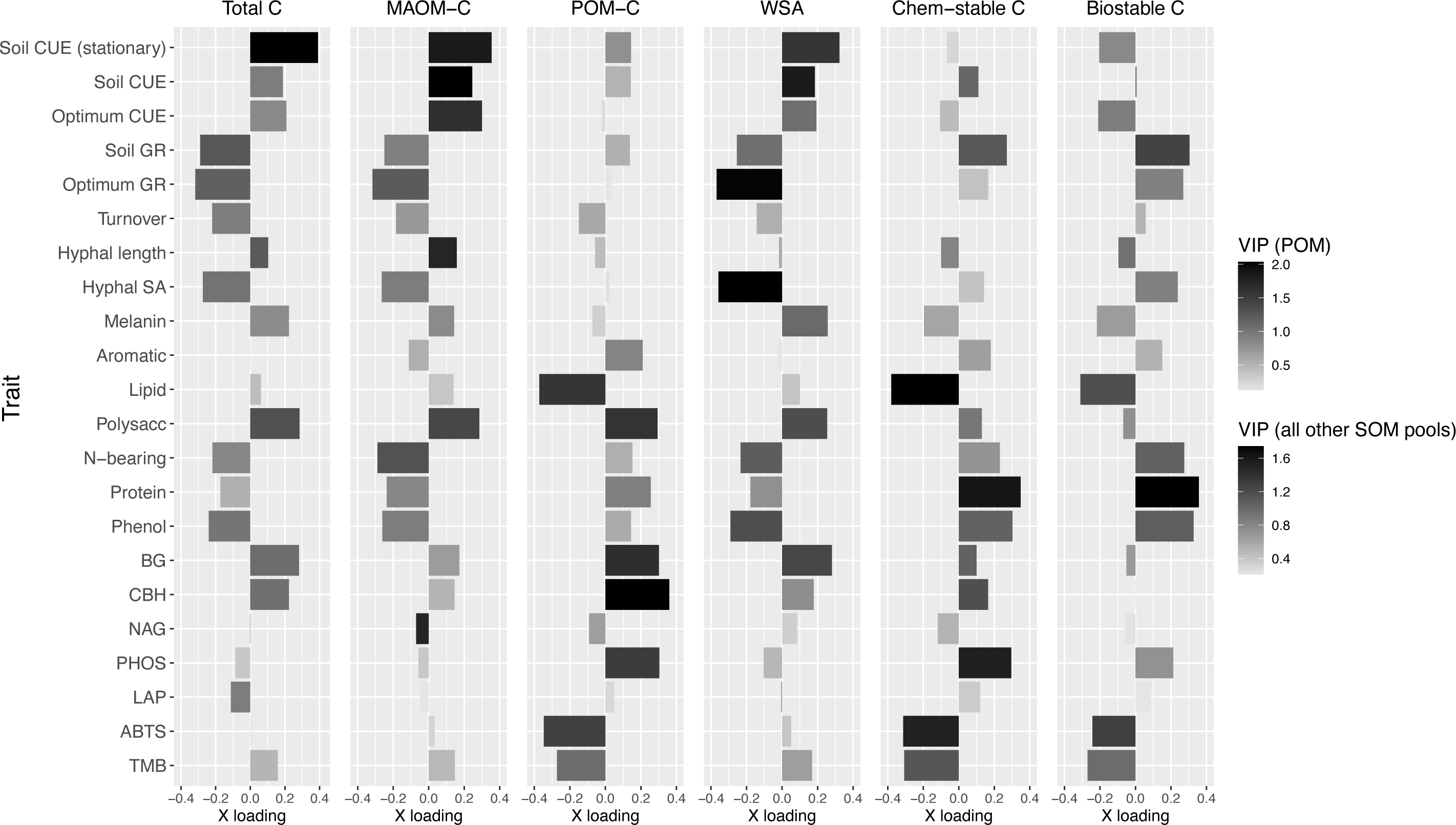
Bar plots of trait loadings on the first (most explanatory) PLSR latent factor for each SOM functional pool, accounting for 50-78% of variation in each pool. From left to right: total C, MAOM-C, POM-C, water-stable aggregates (WSA), chemically stable C, and biologically stable C. This figure corresponds with Fig. 4 in the main text and includes an additional SOM functional pool (POM). X loadings are presented along the x axis, and bar color is shaded to represent variable importance scores (VIP) for each trait variable, indicating the overall importance of each trait to the PLSR model, integrating latent factor 2, and in some cases, factor 3. Full PLSR model outputs are available in Table S1.

**Figure S6.**
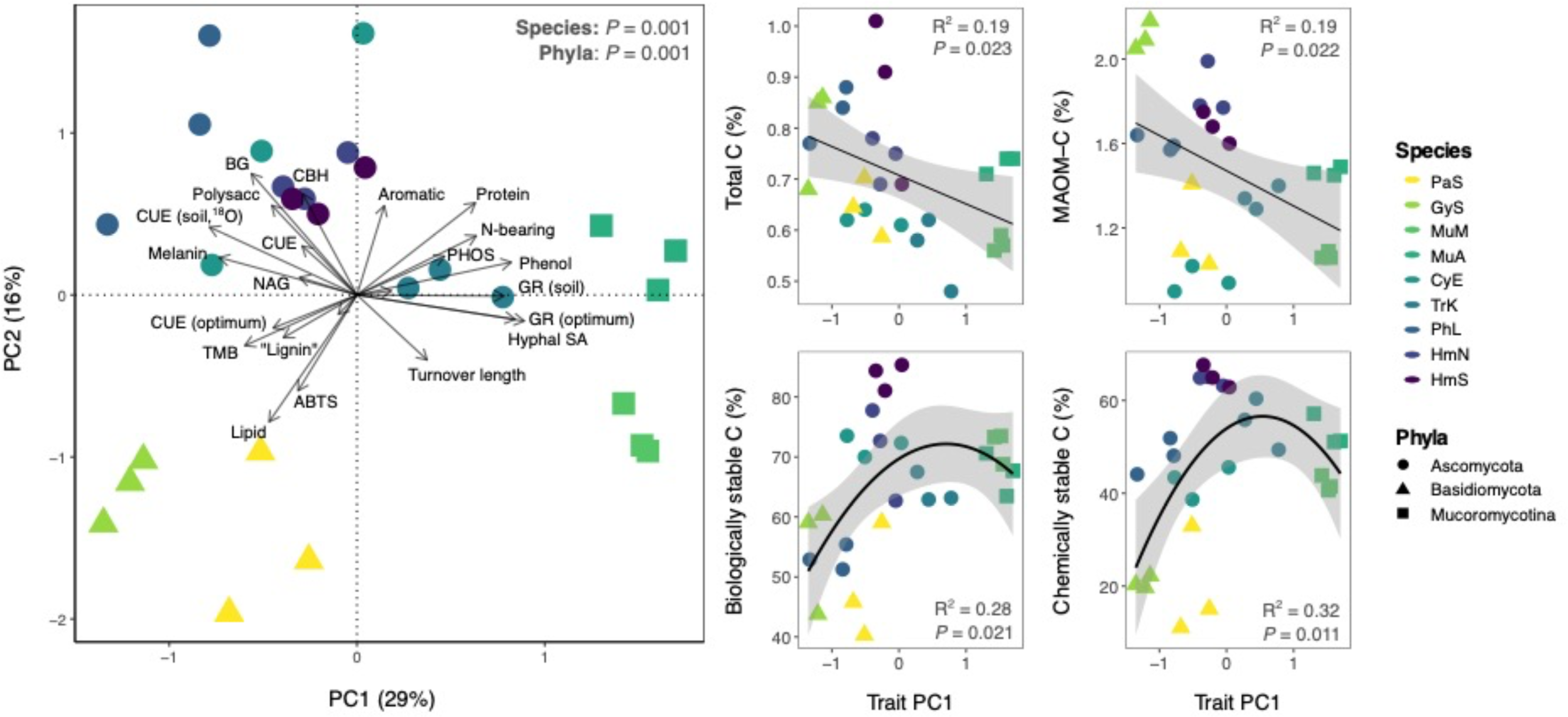
Principal components analysis (PCA) of fungal trait profiles, with the two most explanatory PC axes explaining 45% of variation in fungal trait data. This figure corresponds with Fig. 2 in the main text and includes additional correlation plots for PC axis 1 versus four different SOM functional pools: total C, MAOM-C, biologically stable C and chemically stable C. Sample point color represents fungal species, while point shape represents fungal phyla. P-values and R^2^ are presented for the polynomial regression model with the lowest mean square error for each SOM functional pool.

**Figure S7.**
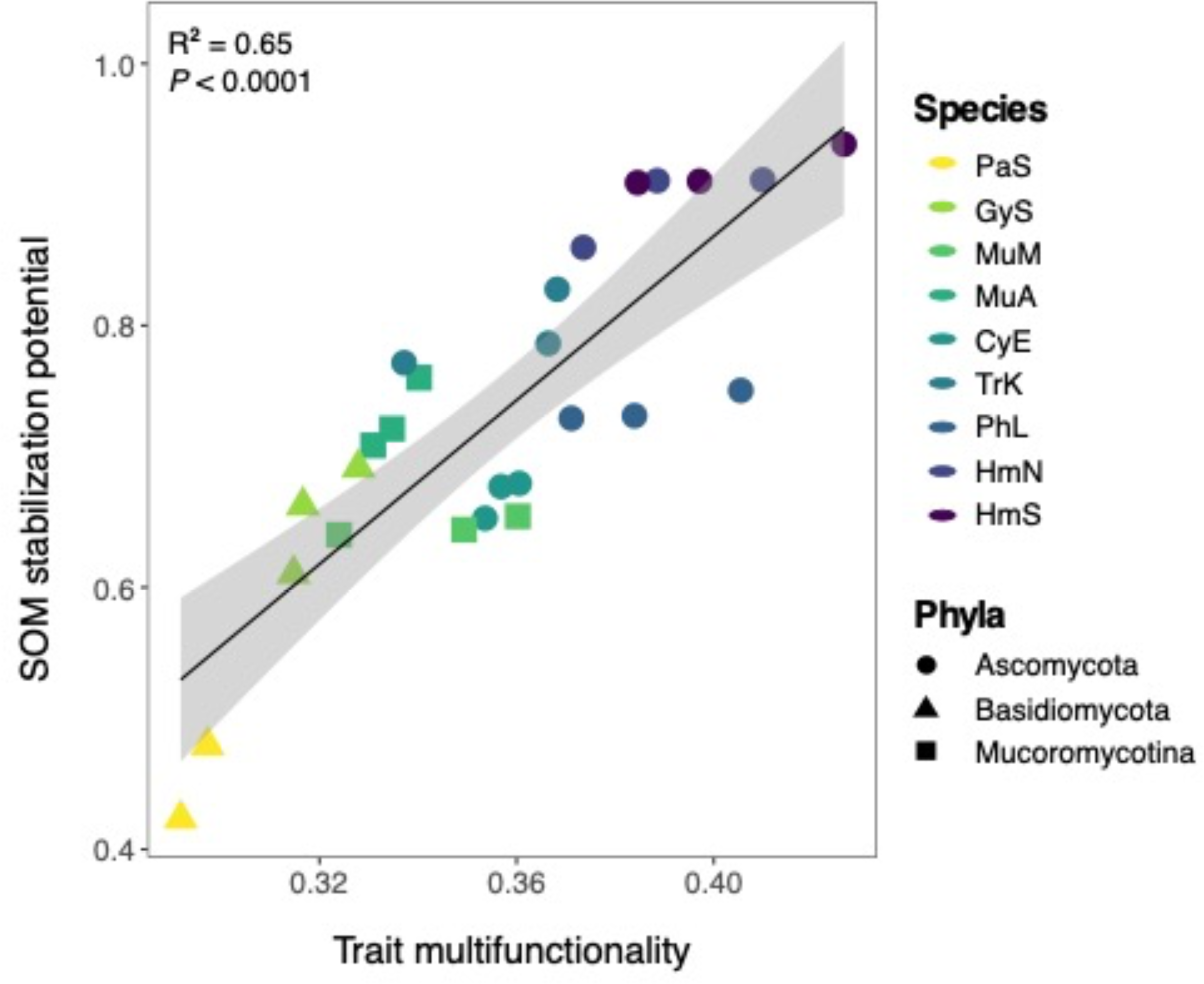
Linear regression between trait multifunctionality and SOM ‘stabilization’ potential (multifunctionality), which included only those SOM functional pools that are putatively stable (MAOM-C, water-stable aggregates, biologically stable C, chemically stable C). Point color represents fungal species, while point shape represents fungal phyla. One PaS replicate with significantly lower trait multifunctionality (∼0.2) is removed from the plot for ease of visualizing differences across other samples, but does not change the interpretation of the results (both *P* < 0.0001). ANOVA results are presented for the model that included all replicates (R^2^ = 0.65; *P* < 0.0001).

**Figure S8.**
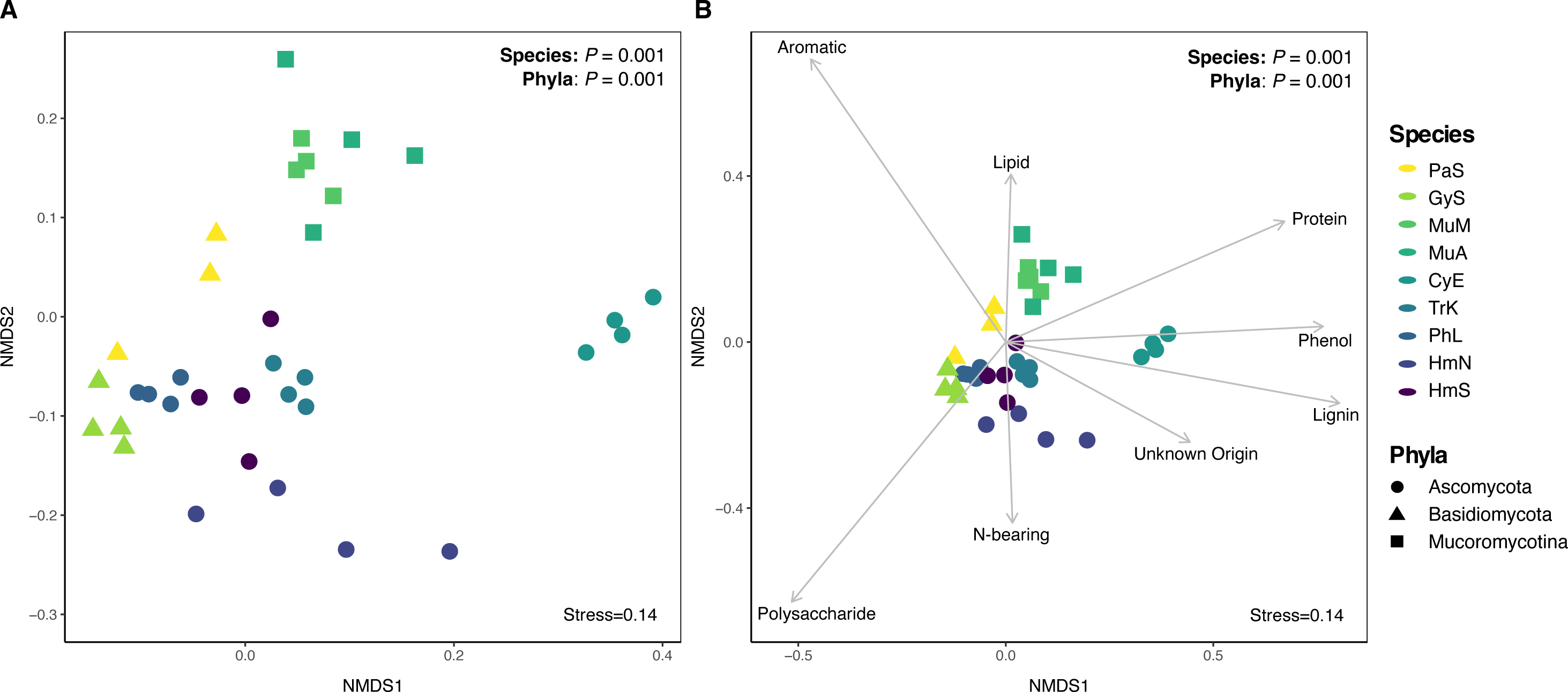
NMDS ordination of MAOM chemical composition (individual compound-level dataset) after long-term incubation with fungal isolates, without environmental vectors (A) or with significant (*P* < 0.05) environmental vectors (B) representing broad chemical compound classes. Version without environmental vectors (A) is included for ease of visualizing differences across samples. Sample point color represents fungal species, while point shape represents fungal phyla. Results of PERMANOVA analyses for both species and phyla are included.

**Figure S9.**
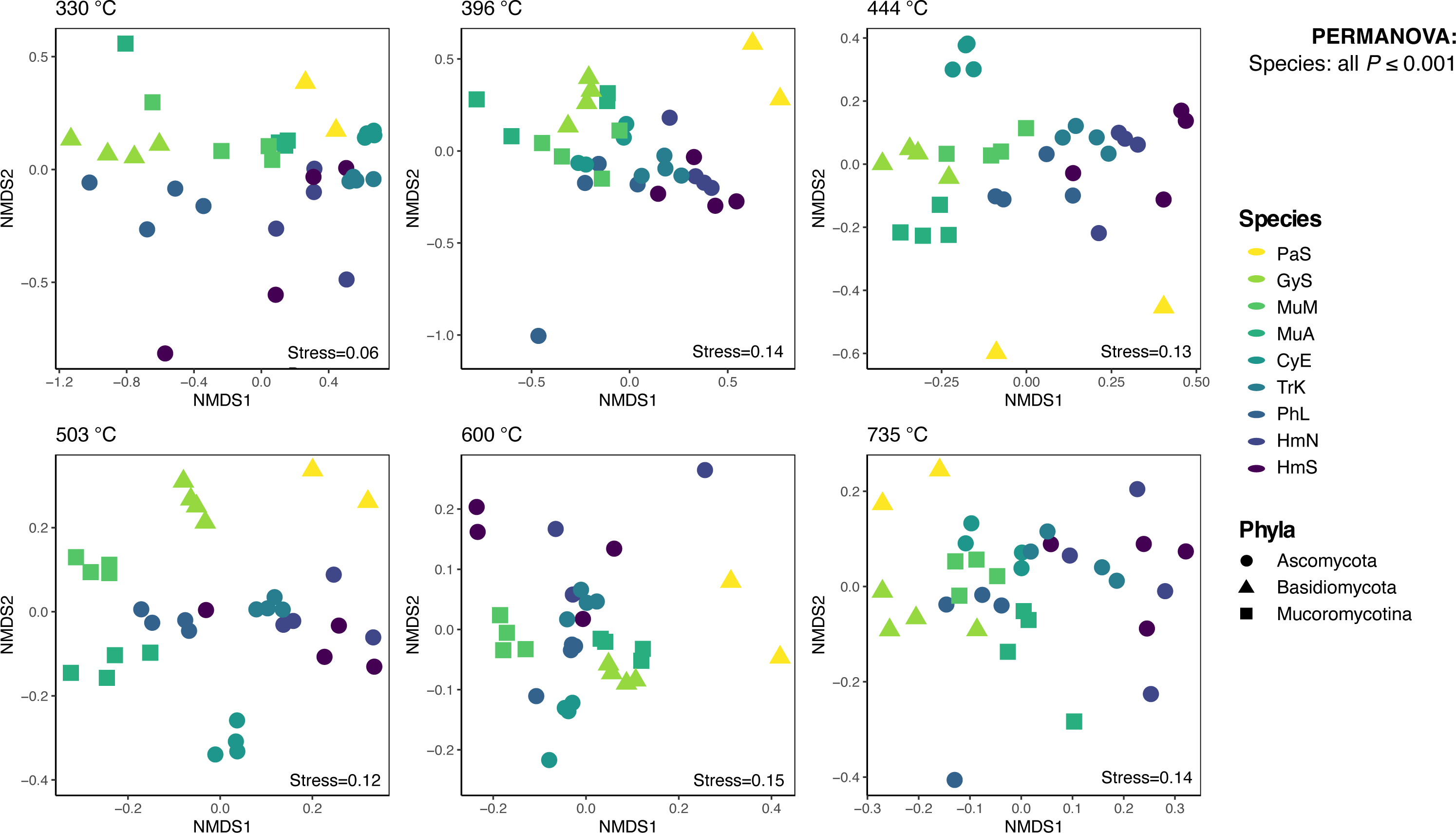
NMDS ordinations of SOM chemical composition after long-term incubation with fungal isolates. Separate panels are shown for each pyrolysis thermal fraction (ramped pyrolysis GC/MS approach), 330°C-735°C. Significant variation in species’ SOM chemistries were observed for each thermal fraction (all *P* ≤ 0.001; PERMANOVA). Point color represents fungal species, while point shape represents fungal phyla.

**Figure S10.**
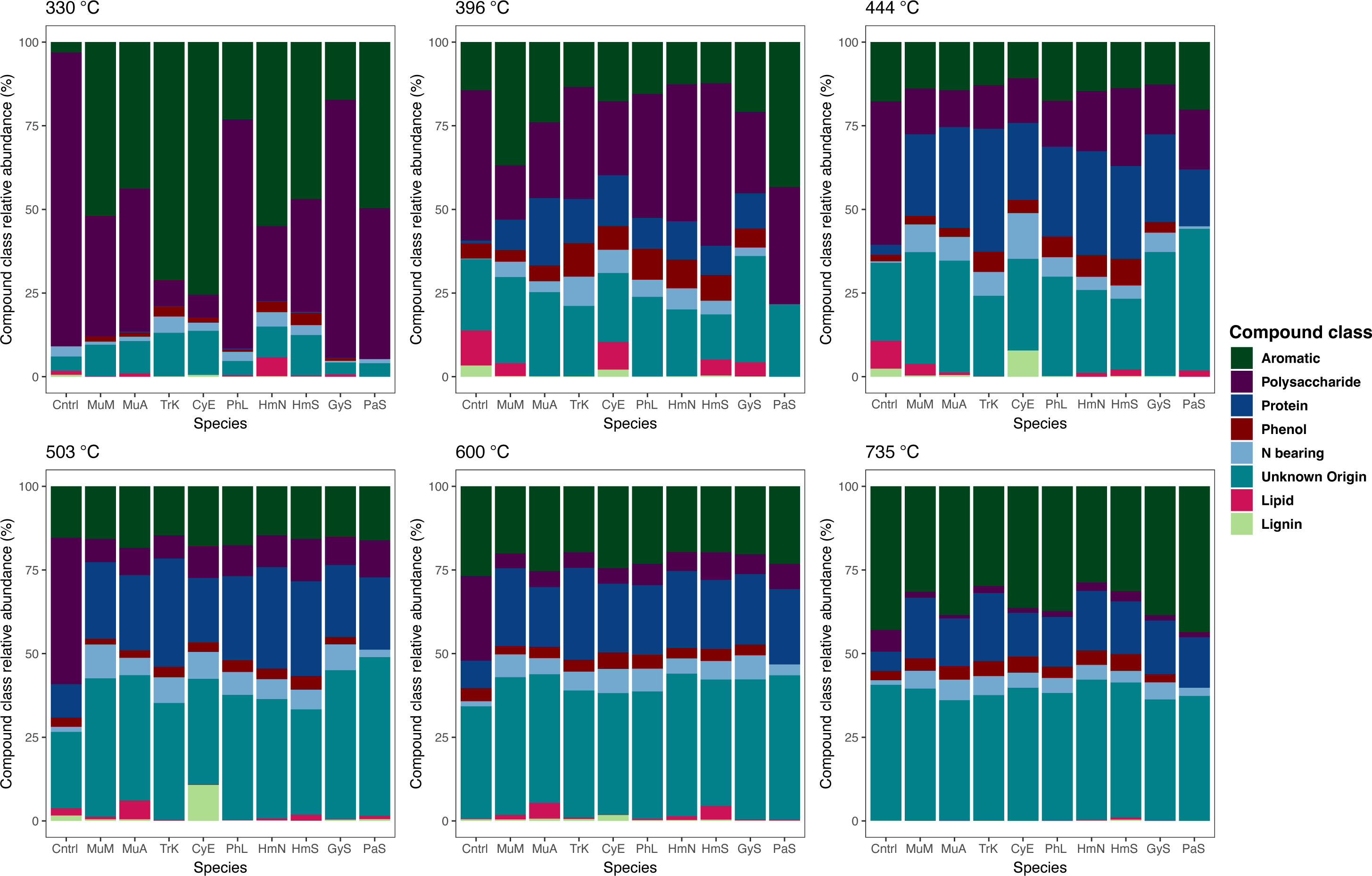
Stacked bar plots showing SOM chemical composition after long-term incubation with fungal isolates, with a separate panel for each pyrolysis thermal fraction (ramped pyrolysis GC/MS approach; 330°C-735°C). Bar segment color represents the relative abundance (%) of broad chemical compound classes (aromatics, polysaccharides, proteins, phenols, N-bearing compounds, lipids, “lignin” and unknown origin). “Lignin” is hypothesized to represent fungal-derived melanins and may have been misidentified by Py-GC/MS analysis of the model soils. This figure corresponds with the NMDS plots presented in Fig. S8.

**Figure S11.**
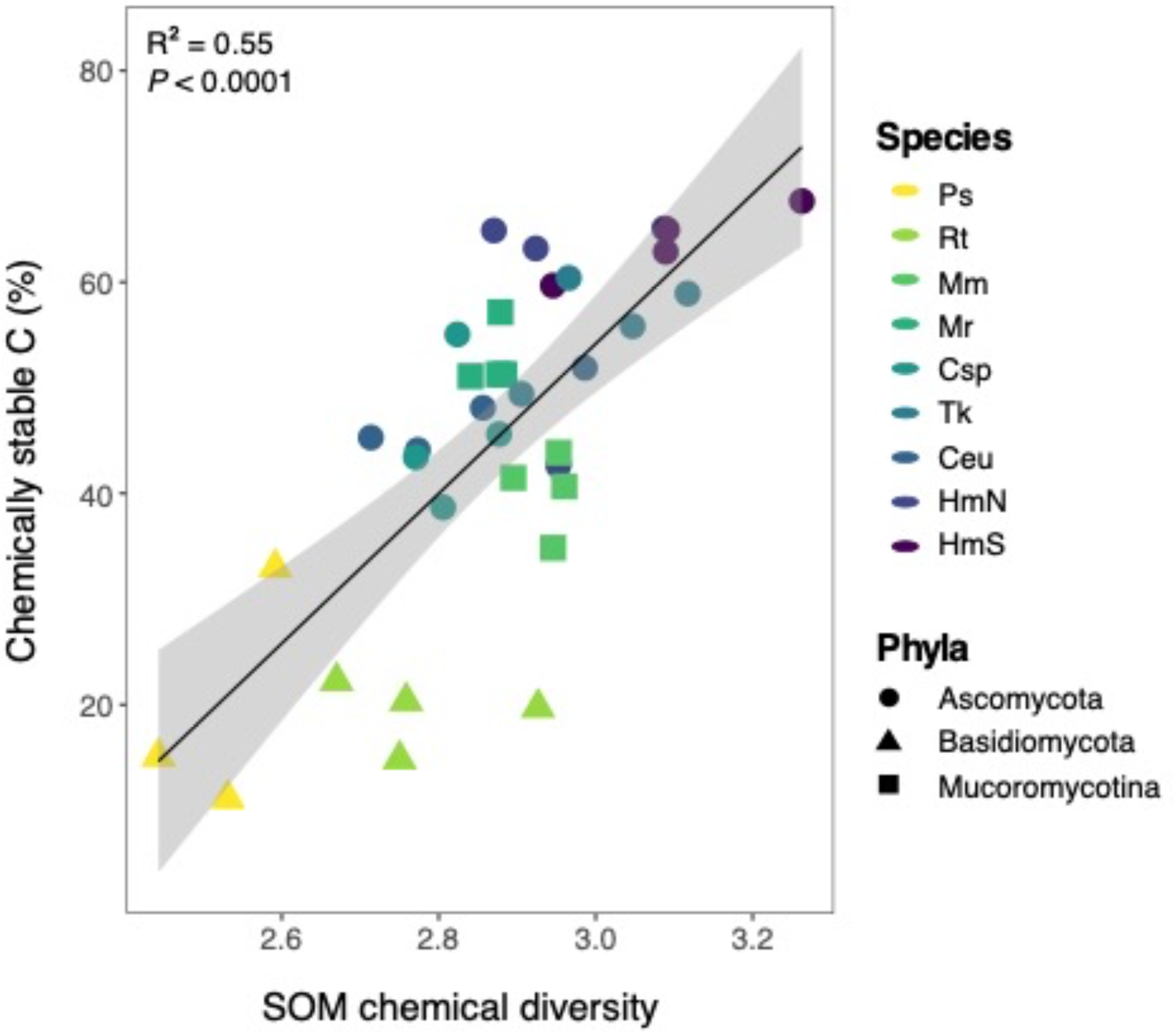
Chemical diversity of thermally stable SOM (735°C) versus the proportion of chemically stable SOM produced by fungal species. Chemical diversity was calculated using the index for Shannon diversity. Sample point color represents fungal species, while point shape represents fungal phyla.

**Figure S12.**
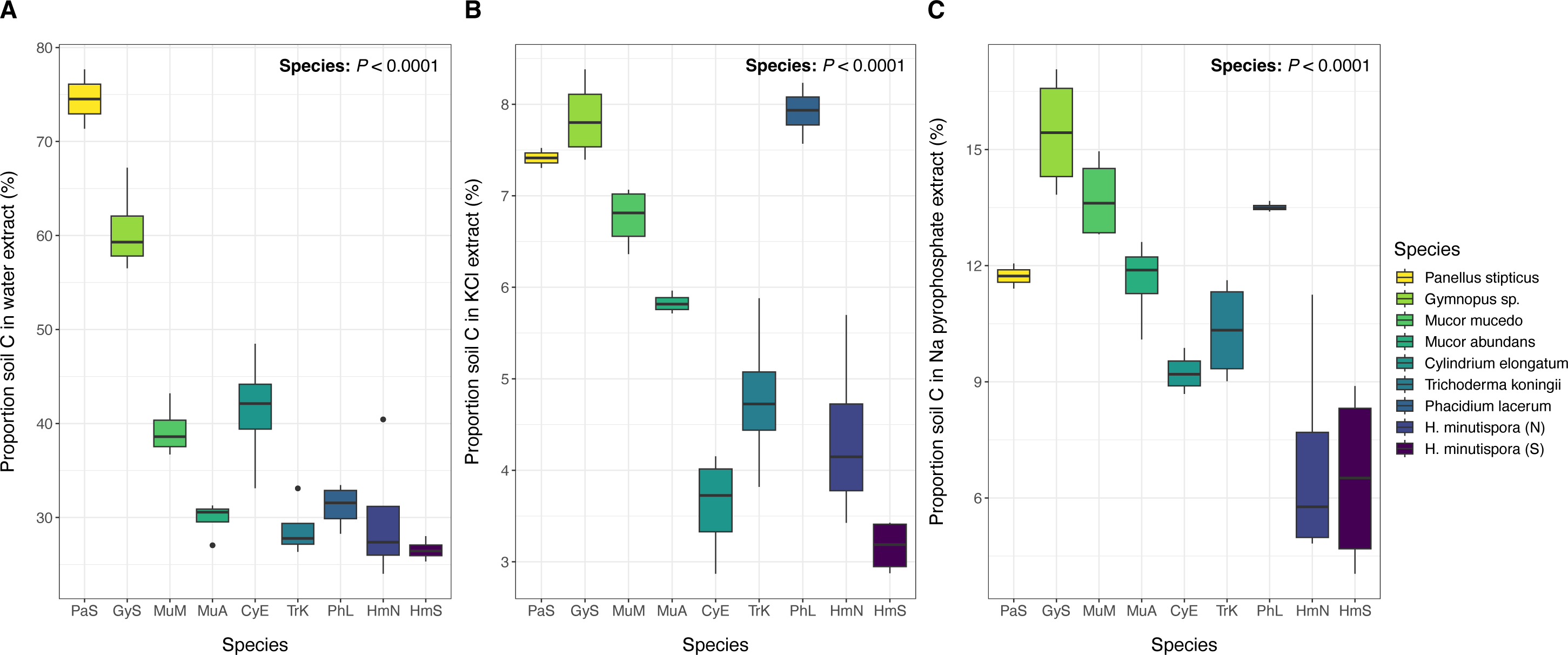
Proportion of fungal-derived SOM-C extracted by (A) water, (B) KCl, or (C) sodium (Na) pyrophosphate during sequential extraction for each fungal isolate. While a substantial portion of soil C was extracted by water, KCl and sodium pyrophosphate for the two Basidiomycota species (PaS, GyS), for many of the Ascomycota (and to a lesser extent the Mucoromycotina) a significant portion of soil C was unable to be extracted by the three extractants and remained in the soil pellet. This remaining fraction of soil C was termed “chemically stable C” (up to 60% for some species; presented in main text).

